# Prenatal exposure to maternal stress drives neurodevelopmental disruptions in the fetal hypothalamus

**DOI:** 10.64898/2026.06.08.730949

**Authors:** Kodee Bao, Matthew Rosin, Jessica M. Rosin

## Abstract

The hypothalamus plays a central role in integrating physiological stressors to maintain homeostasis, yet how fetal neurodevelopment in the hypothalamus is shaped by intrauterine maternal stress exposure remains understudied. This is especially true in the context of sex-divergent mechanisms underlying neurodevelopmental disorders (NDDs), which are increasingly being linked to perturbation of the intrauterine environment. Herein, we utilize a mouse model of prenatal maternal cold stress exposure to study the impacts on neural stem and progenitor cell (NSPC) developmental programs in the fetal hypothalamus. Pregnant mice were exposed to cold stress from embryonic day 11.5 (E11.5) to E15.5 and fetal hypothalamic NSPCs from both male and female embryos were analyzed. Exposure to maternal stress altered fetal hypothalamic neurodevelopment, increasing TUJ1+ neuron number in males, while enhancing neuronal dendritic arborization in females. To define underlying molecular changes, we performed single-cell RNA sequencing of hypothalamic NSPCs. Interestingly, we identified distinct baseline transcriptional profiles between male and female NSPCs. In females, maternal stress upregulated pathways related to GABAergic differentiation and neuronal projection morphogenesis, with these alterations being maintained across more differentiated neuronal populations. Ligand-receptor analysis further indicated that maternal stress alters cell-cell communication within NSPCs, predominantly in females. Together, these findings demonstrate that prenatal maternal stress alters hypothalamic NSPC developmental programs uniquely in each sex, and suggest that disrupted intercellular signaling may contribute to underlying sex differences in social behaviors previously reported for this model (Rosin et al., 2021).

**SIGNIFICANCE STATEMENT:** Prenatal stress is a known risk factor for NDDs, but how it shapes early brain development in a sex-specific manner remains understudied. Here, we examined how maternal stress influences NSPCs in the hypothalamus, a brain region critical for regulating the stress response and homeostasis. Using mice as a model system, we found that maternal stress alters how fetal NSPCs develop into neurons uniquely in each sex. Molecular analyses demonstrate that maternal stress alters female NSPCs transcriptional programs and cell-to-cell communication. This work advances our understanding of how prenatal maternal stress drives sex differences in neurodevelopmental programming and may help to begin to explain sex-biased vulnerabilities to NDDs.

## INTRODUCTION

During embryogenesis, the brain develops from neural stem and progenitor cells (NSPCs), which self-renew and differentiate into neurons, oligodendrocytes, and astrocytes, each defined by distinct transcriptional profiles (Kim et al., 2020, 2025; Fong et al., 2023). In mice, hypothalamic neurogenesis occurs predominantly between embryonic day 10.5 (E10.5) and E16.5, while gliogenesis initiates around E13.5 (Marsters et al., 2016; Fong et al., 2023). Beyond intrinsic transcriptional regulation, accumulating evidence shows that neurodevelopment is also shaped by interactions between NSPCs and microglia (Antony et al., 2011; Cunningham et al., 2013). Further supporting this interplay, high-resolution imaging demonstrates close interactions between microglia and NSPCs within the developing hypothalamus, including upregulation of cytokine-receptor pathways associated with stress signaling (Rosin et al., 2021).

A healthy intrauterine environment is critically important for neurodevelopment, with disruptions during this window altering NSPC behavior and circuit formation. A wide range of adversities can influence the intrauterine environment and neurodevelopment, including adverse psychological states (e.g., anxiety, depression, etc.), physiological challenges (e.g., infection, inflammation, etc.), and environmental exposures (e.g., pollution, toxicants, etc.), which are often collectively conceptualized as prenatal maternal stress (Hossain and Westerlund Triche, 2007; Buss et al., 2011; Bergh et al., 2018; Mareckova et al., 2020; Yu et al., 2023). Despite their diverse origins, these factors are consistently associated with altered fetal brain development and increased risk of neurodevelopmental disorders (NDDs), as supported by epidemiological studies linking prenatal maternal stress to changes in offspring brain structure and mental health outcomes (Buss et al., 2011; Favaro et al., 2015; Bergh et al., 2018; Marečková et al., 2019; Mareckova et al., 2020; Goldstein et al., 2021; Lu et al., 2022; Yu et al., 2023).

Maternal immune activation has been proposed as a key pathway linking prenatal maternal stress to altered fetal neurodevelopment, whereby maternal stress triggers changes in corticosterone levels and/or inflammatory signaling that can propagate to the fetal brain, with fetal microglia acting as key mediators of altered neurodevelopment (Bolton et al., 2014; Zhao et al., 2014; Rosin et al., 2021; Shimizu et al., 2023; Woods et al., 2023; Yu et al., 2023). Supporting this, a recent study examining maternal cytokine levels in a human prenatal cohort found that imbalances between pro- and anti-inflammatory cytokines during pregnancy were associated with sex-specific changes in adult brain circuitry, particularly within the hypothalamus (Goldstein et al., 2021). Relevant to the current study, cold exposure represents a physiologically relevant environmental stressor, driving alterations to maternal, placental, and embryonic signaling that are characteristic of the physiological changes seen in maternal stress models (Lian et al., 2017; Rosin et al., 2021). Indeed, cold exposure has been shown to elevate maternal corticosterone levels and induce changes in inflammatory signaling in the placenta (Lian et al., 2017). Moreover, prenatal exposure to cold stress highlights a male-specific sensitivity of microglia to maternal stress, as intermittent cold exposure increases NSPC-residing microglia numbers, elevates the secretion of pro-inflammatory cytokines and chemokines, decreases oxytocin neurons in the paraventricular nucleus (PVN), and leads to microglia-dependent social deficits in offspring, but only in males (Rosin et al., 2021). Importantly, female offspring exposed to prenatal maternal cold stress during gestation also develop social deficits; however, this is independent of microglia (Rosin et al., 2021), suggesting that neurodevelopmental disruptions underlying social deficits seen in offspring later in life likely differ between males and females.

Despite these findings, it remains unclear how exposure to prenatal maternal cold stress alters fetal hypothalamic NSPC developmental programs and whether these effects differ between males and females. Here, we utilize neurosphere and differentiation assays to investigate how maternal stress affects fetal hypothalamic NSPC proliferation, differentiation, stem-like potential, and lineage trajectories, showing impacts on neuronal numbers and branching complexity. We also employ single-cell RNA sequencing (scRNAseq) to examine how maternal stress impacts the transcriptomic profile of fetal hypothalamic NSPCs and identify upregulation of pathways involved in GABAergic differentiation and neuronal projection morphogenesis uniquely in females. Our findings demonstrate that maternal stress alters fetal NSPCs through sex-divergent mechanisms, providing insight into how early-life insults may contribute to sex differences observed in many NDDs.

## RESULTS

### Prenatal maternal cold stress exposure does not disrupt the proliferative capacity or stem-like potential of fetal hypothalamic NSPCs

Microglia reside in close proximity to hypothalamic NSPCs during embryogenesis and exhibit sex differences in their responsiveness to prenatal maternal cold stress exposure (Rosin et al., 2021). Although maternal cold stress exposure during pregnancy has been shown to disrupt oxytocin neuron numbers in the fetal PVN and social behaviors in offspring postnatally uniquely in each sex, the upstream developmental mechanisms driving these outcomes remain unclear (Rosin et al., 2021). These findings suggest that stress-induced alterations in microglial signaling may influence hypothalamic neurodevelopment by acting at the level of NSPCs. At the same time, NSPCs themselves may be directly responsive to prenatal stress; however, the extent to which prenatal maternal cold stress exposure alters NSPC behaviors has not yet been investigated. Accordingly, we examined how prenatal maternal cold stress impacts hypothalamic NSPC proliferation and stem-like potential using a neurosphere assay (Reynolds and Weiss, 1992; Nesan et al., 2020; Rosin et al., 2021). Briefly, hypothalamic NSPCs were isolated from embryonic day 15.5 (E15.5) control and prenatal maternal cold stress-exposed male and female embryos and cultured in 24-well plates for 10 days to form primary neurospheres (Figure 1A).

**Figure 1.**
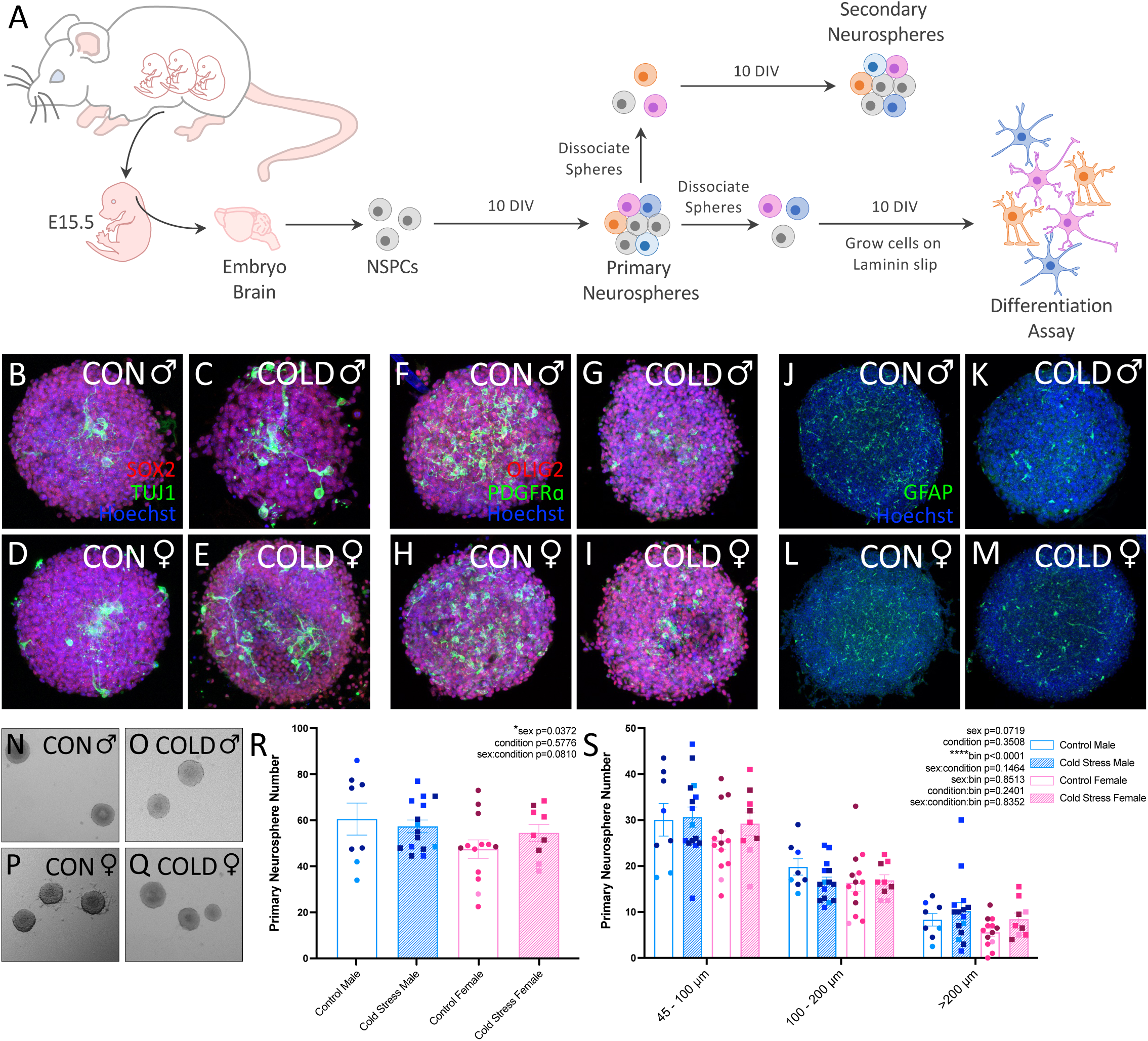
Maternal cold stress exposure does not impact the proliferative capacity of E15.5 NSPCs. (A) Schematic diagram illustrating the neurosphere assay workflow. Days *in vitro* (DIV). (B-E) Primary neurospheres stained with SOX2, TUJ1, and Hoechst. (F-I) Primary neurospheres stained with OLIG2, PDGFRα, and Hoechst. (J-M) Primary neurospheres stained with GFAP and Hoechst. (N-Q) Representative images of primary neurospheres generated from E15.5 male and female NSPCs. (R) Quantification of the number of primary neurospheres generated from E15.5 control male (n=8 embryos; N=3 litters), control female (n=13 embryos; N=3 litters), cold stress male (n=15 embryos; N=3 litters), and cold stress female (n=9 embryos; N=3 litters) NSPCs. (S) Quantification of the size distribution of primary neurospheres generated from E15.5 control male, control female, cold stress male, and cold stress female NSPCs, which were binned by sphere sizes between 45-100µm, 100-200µm or >200µm. Counts represent means ± SEM and were analyzed using a mixed effects model with litter incorporated as a random effect, where post hoc tests were evaluated using a Bonferroni correction. (R, S) Each dot represents primary neurospheres derived from a single, independent embryo. Dots of same color represent embryos from the same litter, while dots of different color represent embryos from different litters (N=3 independent litters).

E15.5 was selected as the time-point for assessment as this is a critical period of neurodevelopment in the fetal hypothalamus whereby microglia are lining the NSPC niche and commitment of NSPCs is transitioning from neurogenic to gliogenic programs (Marsters et al., 2016; Rosin et al., 2021; Fong et al., 2023). Immunofluorescence staining confirmed that primary neurospheres contained sex-determining region Y (SRY)-box 2 transcription factor (SOX2+) neural progenitors and class III β-tubulin (TUJ1+) neurons (Figures 1B-E), oligodendrocyte transcription factor 2 (OLIG2+) and platelet-derived growth factor receptor alpha (PDGFRα+) oligodendrocyte precursor cells (OPCs)/oligodendrocytes (Figures 1F-I), and glial fibrillary acidic protein (GFAP+) astrocytes (Figures 1J-M). Quantification of primary neurospheres derived from control and prenatal maternal cold stress-exposed male and female hypothalamic NSPCs (Figures 1N-Q) showed no impact of prenatal maternal cold stress exposure on the number of neurospheres formed (Figure 1R; condition F_(1,39)_=0.3154, p=0.5776); however, there was a main effect of sex (Figure 1R; sex F_(1,39)_=0.4.654, p=0.0372). While an overall sex difference in the number of primary neurospheres was detected, it could not be attributed to any specific group differences (Figure 1R, Table S1; control male vs. female p=0.0710; prenatal maternal cold stress male vs. female p=1). Quantification of primary neurospheres ranging in size from 45µm to >200µm in diameter showed no impact of prenatal maternal cold stress exposure on male or female neurosphere size (Figure 1S; condition F_(1,39)_=0.8919, p=0.3508), irrespective of whether the spheres were 45-100µm, 100-200µm or >200µm in diameter (Table S1). Interestingly, there was a main effect of neurosphere size (Figure 1S; bin F_(2,82)_=143.6, p<0.0001), with the majority of male and female neurospheres having a diameter of 45-100µm and fewer male and female neurospheres having a diameter >200µm (Table S1).

Next, we examined how prenatal maternal cold stress exposure impacts the stem-like potential or “stemness” of hypothalamic NSPCs by generating secondary neurospheres. Immunofluorescence staining confirmed that secondary neurospheres contained SOX2+ neural progenitors (Figures S1A-D), OLIG2+ and PDGFRα+ OPCs/oligodendrocytes (Figures S1E-H), and GFAP+ astrocytes (Figures S1I-L); however, TUJ1+ neurons were not observed. Quantification of secondary neurospheres derived from the dissociation of control and prenatal cold stress-exposed male and female primary neurospheres (Figures S1M-P) showed no impact of prenatal maternal cold stress exposure on the number of secondary neurospheres formed (Figure S1Q; condition F_(1,33)_=1.012, p=0.3217). Similarly, quantification of secondary neurospheres ranging in size from 45µm to >200µm in diameter showed no impact of prenatal maternal cold stress exposure on male or female neurosphere size (Figure S1R; condition F_(1,33)_=1.012, p=0.3217), irrespective of whether the spheres were 45-100µm, 100-200µm or >200µm in diameter (Table S1).

Interestingly, secondary neurospheres showed a main effect of size (Figure S1R; bin F_(2,70)_=52.83, p<0.0001), but this time with the majority of male and female neurospheres having a diameter >200µm and fewer male and female neurospheres having a diameter between 45-100µm (Table S1).

Together, these findings suggest that prenatal maternal cold stress exposure does not disrupt the proliferative capacity or stem-like potential of male or female hypothalamic NSPCs at this particular time-point during gestation when using an *in vitro* neurosphere assay.

### Exposure to prenatal maternal cold stress alters NSPC differentiation and neuronal morphology

Given the capacity of NSPCs to differentiate into neurons and glia, we used a differentiation assay to examine the effects of prenatal maternal cold stress exposure on lineage outcomes by quantifying neuron, OPC, oligodendrocyte, and astrocyte numbers (Kim et al., 2020, 2025; Fong et al., 2023). To achieve this, primary neurospheres were mechanically dissociated and plated on Laminin slips for 10 days and allowed to differentiate (Figure 1A). Five unique, non-overlapping fields of view were imaged per slip in a consistent pattern to ensure representative sampling of cellular distribution, and the number of each differentiated cell-type (e.g., neuron, OPC, oligodendrocyte or astrocyte) was quantified as a proportion of the total number of Hoechst+ cells observed in the field of view. Proportions across the five fields of view were averaged to give one representative value for the entire slip, where each slip was generated from primary neurospheres obtained from a single, independent embryo. Interestingly, when comparing control and prenatal maternal cold stress male and female cells on slips stained with TUJ1 (Figures 2A-D), we observed an impact of prenatal maternal cold stress on the proportion of TUJ1+ neuron numbers (Figure 2E; condition F_(1,18)_=11.96, p=0.0028). Specifically, a significant increase in the proportion of TUJ1+ neurons was observed in prenatal maternal cold stress males compared to control males (Figure 2E; p=0.0229), which was not seen in females (Figure 2E, Table S2; control female vs. prenatal maternal cold stress female p=0.8002).

**Figure 2.**
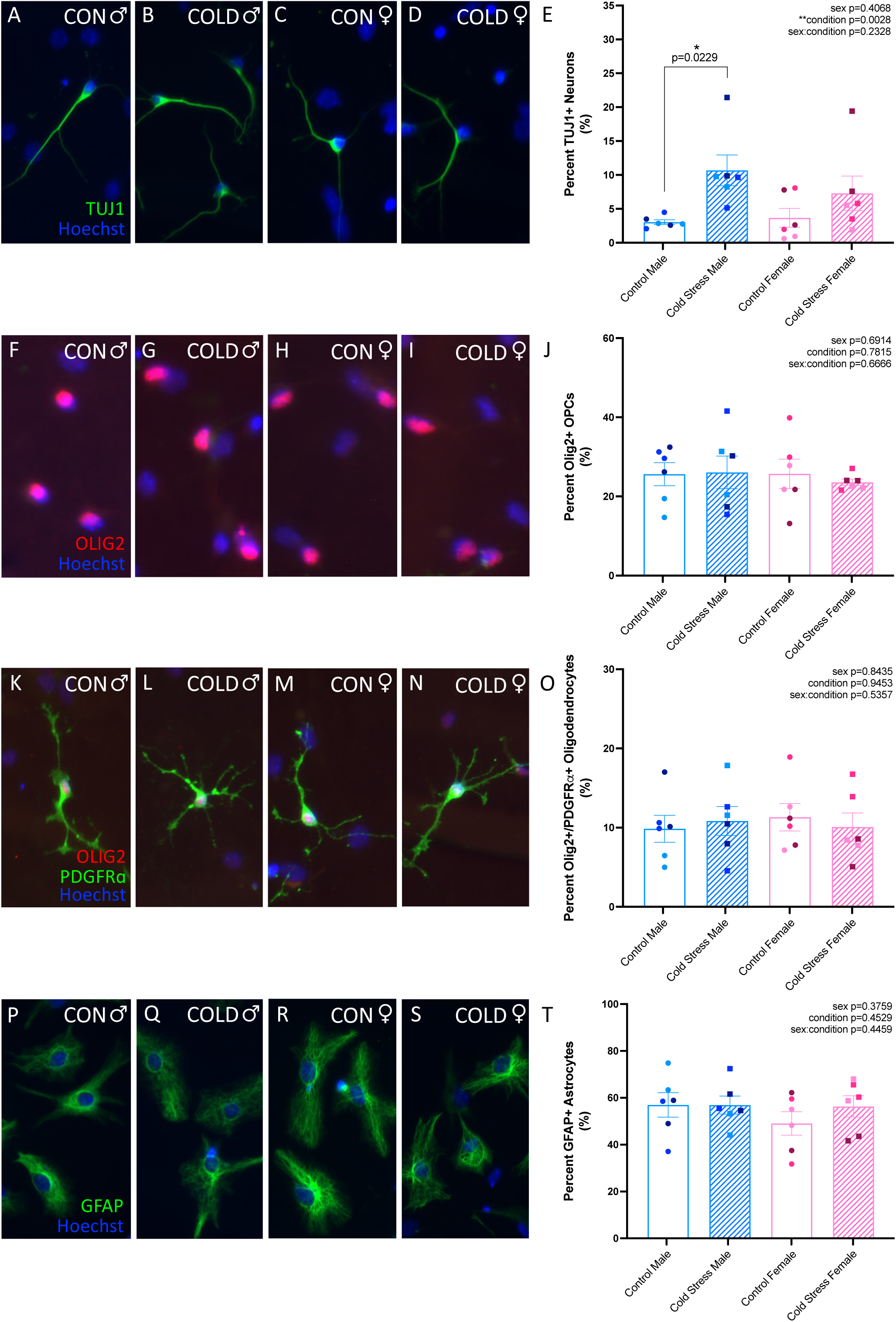
Maternal cold stress exposure significantly increases the number of TUJ1+ neurons in male embryos without impacting glial cell differentiation in either sex. (A-D) Representative images of differentiated neurons derived from male and female primary neurospheres that were dissociated and plated on laminin-coated glass slips and stained with TUJ1 and Hoechst. (E) Percent of Hoechst+ control male and female or cold stress male and female cells on laminin-coated glass slips that were TUJ1+ neurons. (F-I) Representative images of differentiated OPCs derived from male and female primary neurospheres that were dissociated and plated on laminin-coated glass slips and stained with OLIG2 and Hoechst. (J) Percent of Hoechst+ control male and female or cold stress male and female cells on laminin-coated glass slips that were OLIG2+ OPCs. (K-N) Representative images of differentiated oligodendrocytes derived from male and female primary neurospheres that were dissociated and plated on laminin-coated glass slips and stained with OLIG2, PDGFRα, and Hoechst. (O) Percent of Hoechst+ control male and female or cold stress male and female cells on laminin-coated glass slips that were OLIG2+/PDGFRα+ double-positive oligodendrocytes. (P-S) Representative images of differentiated astrocytes derived from male and female primary neurospheres that were dissociated and plated on laminin-coated glass slips and stained with GFAP and Hoechst. (T) Percent of Hoechst+ control male and female or cold stress male and female cells on laminin-coated glass slips that were GFAP+ astrocytes. (E, J, O, T) Each dot represents a glass slip of differentiated cells generated from primary neurospheres that were derived from a single, independent embryo (n=6 embryos). Dots of same color represent embryos from the same litter, while dots of different color represent embryos from different litters (N=3 independent litters). Counts represent means ± SEM and were analyzed using a mixed effects model with litter incorporated as a random effect, where post hoc tests were evaluated using a Bonferroni correction.

Next, we examined whether prenatal maternal cold stress exposure altered OPC or oligodendrocyte lineages. Interestingly, no difference in the proportion of OPCs was observed between control and prenatal maternal cold stress conditions in either male- or female-derived cell cultures that were plated on slips and stained with OLIG2 (Figures 2F-J; condition F_(1,18)_=0.0793, p=0.7815). Moreover, quantification of the proportion of OLIG2+/PDGFRα+ double-positive oligodendrocytes also showed no differences between control and prenatal maternal cold stress conditions for both males and females (Figures 2K-O; condition F_(1,20)_=0.0048, p=0.9453). Similar to what was observed for OPC and oligodendrocyte glial lineages, quantification of the proportion of GFAP+ astrocytes showed no differences between control and prenatal maternal cold stress conditions for both males and females (Figures 2P-T; condition F_(1,20)_=0.5860, p=0.4529).

Based on these findings, we decided to examine whether prenatal maternal cold stress exposure impacts neuronal morphology by performing morphological analysis (Schindelin et al., 2012). Although formal classifications of neuronal morphology have been established primarily in the cortex, similar morphologies were observed in our hypothalamic neuronal cultures (Mason and Bern, 1977; Kriegstein and Dichter, 1983; Mir et al., 2020). Specifically, we could observe three major neuronal morphologies: bipolar, multipolar (stellate-like), as well as neurons exhibiting a polarized morphology characterized by a single, prominent axonal process and a distinct arrangement of shorter dendritic processes (Mir et al., 2020). Consistent with established definitions, bipolar neurons exhibited two opposing processes, while multipolar neurons displayed multiple dendritic branches of relatively uniform morphology. In contrast, the polarized neurons were readily distinguished by their clear axon-dendrite polarity and constituted the largest population of neurons observed within our hypothalamic cultures. Given their clear morphological features and relative abundance, subsequent analyses focused solely on these polarized neurons. Five neurons were imaged per slip, where each slip was generated from cells obtained from a single, independent embryo. For morphological analyses, the number of primary, secondary, and tertiary dendrites was quantified for each neuron (Figure S2) and averaged across the five neurons imaged in order to obtain a representative value per slip/embryo. Quantitative analysis of dendritic morphology for control and prenatal maternal cold stress male and female neurons (Figures 3A-D) revealed no impact of prenatal maternal cold stress on the total number of dendrites (Figure 3E; condition F_(1,20)_= 2.849, p=0.1069) or cumulative dendritic length (Figure 3F; condition F_(1,20)_= 1.308, p=0.2662). Interestingly, a main effect of sex was observed for both the total number of dendrites (Figure 3E; sex F_(1,20)_=4.804, p=0.0404) and cumulative dendritic length (Figure 3F; sex F_(1,20)_=8.481, p=0.0086). Although an overall sex difference in the total number of dendrites was detected, it could not be attributed to any specific group difference (Figure 3E, Table S3). In contrast, a significant difference between control male and females was observed when analyzing cumulative dendritic length (Figure 3F; p=0.0206).

**Figure 3.**
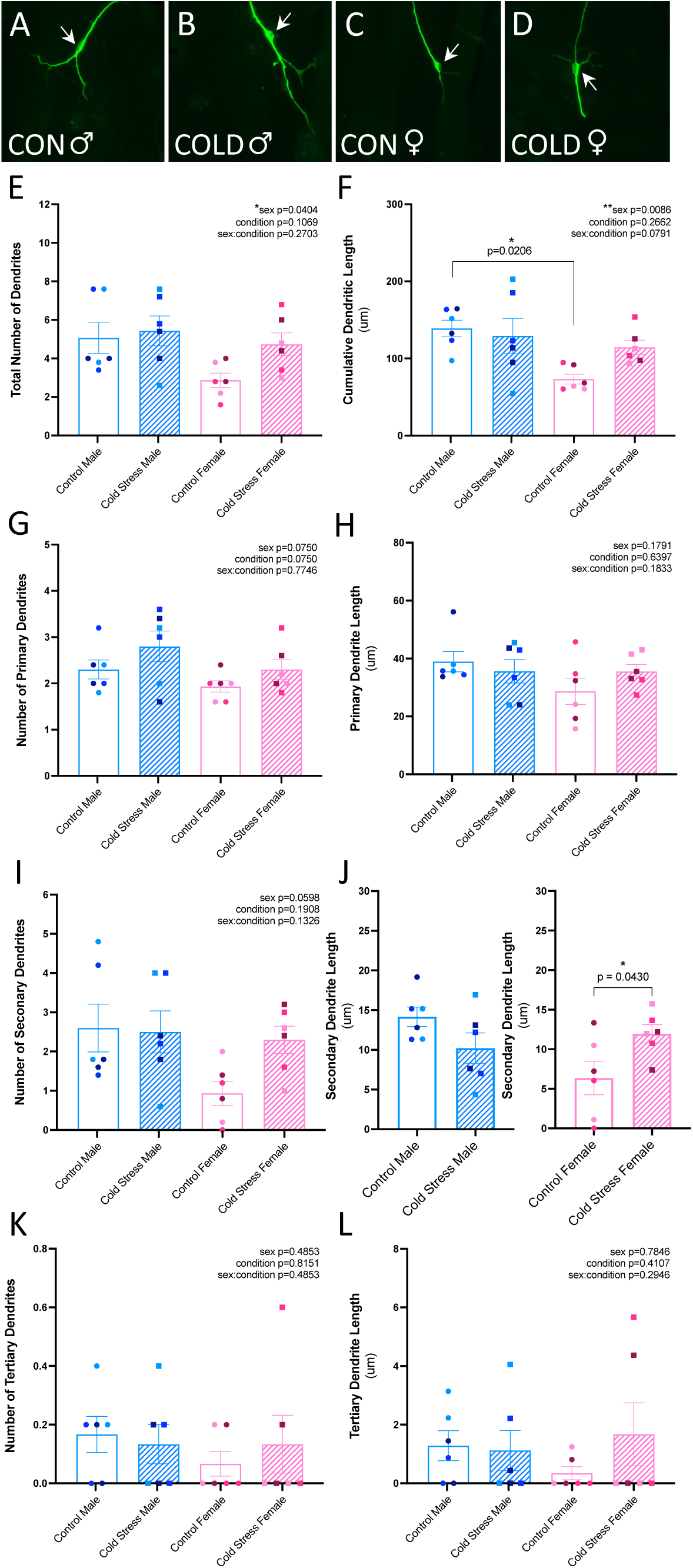
Maternal cold stress exposure significantly increases dendritic arborization in female embryos. (A-D) Representative images of the soma and dendrites of differentiated neurons derived from male and female primary neurospheres that were dissociated and plated on laminin-coated glass slips and stained with TUJ1. Arrows mark the cell soma. (E) Total number of dendrites (primary, secondary, and tertiary dendrites), comparing control and cold stress males and females. (F) Cumulative dendritic length (primary, secondary, and tertiary dendrites), comparing control and cold stress males and females. (G) Number of primary dendrites, comparing control and cold stress males and females. (H) Primary dendrite length, comparing control and cold stress males and females. (I) Number of secondary dendrites, comparing control and cold stress males and females. (J) Secondary dendrite length, comparing control and cold stress males (left) or control and cold stress females (right). (K) Number of tertiary dendrites, comparing control and cold stress males and females. (L) Tertiary dendrite length, comparing control and cold stress males and females. (E-L) Each dot represents a glass slip of differentiated cells generated from primary neurospheres that were derived from a single, independent embryo (n=6 embryos). Dots of same color represent embryos from the same litter, while dots of different color represent embryos from different litters (N=3 independent litters). Counts represent means ± SEM and were analyzed using a mixed effects model with litter incorporated as a random effect, where post hoc tests were evaluated using a Bonferroni correction.

Further analysis of primary dendritic number (Figure 3G; condition F_(1,18)_=3.570, p=0.0750) and length (Figure 3H; condition F_(1,20)_=0.2259, p=0.6397), secondary dendritic number (Figure 3I; condition F_(1,20)_=1.833, p=0.1908), and tertiary dendritic number (Figure 3K; condition F_(1,20)_=0.0562, p=0.8151) and length (Figure 3L; condition F_(1,20)_=0.7059, p=0.4107) indicated that prenatal maternal cold stress exposure did not impact these morphological measurements.

However, as we identified an interaction between prenatal maternal cold stress and sex when evaluating secondary dendritic length (Table S3; sex x condition interaction F_(1,20)_=8.313, p=0.0092), which suggests that males and females show sex-dependent responses to prenatal maternal cold stress exposure, we stratified by sex to determine the magnitude of the response to prenatal maternal cold stress within each sex (Figure 3J). While prenatal maternal cold stress did not result in a significant difference in secondary dendritic length in males (Figure 3J; p=0.1170), it was found to drive a significant increase in secondary dendritic length in females (Figure 3J; p=0.0430).

Collectively, these findings suggest that prenatal maternal cold stress exposure drives changes in the neuronal lineage, with an increased proportion of neurons seen only in males, while the proportion of glial lineages such as OPCs, oligodendrocytes, and astrocytes appear comparable within each sex and treatment condition. Analysis of neuronal morphology also highlights altered dendritic arborization, but only in females.

### Single-cell transcriptomic profiling of fetal hypothalamic neural stem, progenitor, and precursor cells highlights six clusters with unique neuronal or glial trajectories

To explore the transcriptomic profile of NSPCs residing adjacent to and nearby microglia within the developing fetal hypothalamus, and any underlying molecular changes in response to prenatal maternal cold stress exposure, we utilized scRNAseq. Specifically, we micro-dissected and collected the ventricular zone from the E15.5 hypothalamus of both male and female control and prenatal maternal cold stress-exposed embryos and dissociated this tissue into a single-cell suspension (Figure S3A). Sequencing of the pooled libraries yielded high-quality transcriptomic data from E15.5 control and cold stress samples. A total of ∼15,556 cells were recovered from control samples and ∼17,961 cells from cold stress samples. The mean sequencing depth was 24,512 reads per cell for controls and 19,632 reads per cell for cold stress samples, with raw sequencing metrics summarized in Figure S3B.

Downstream filtering and subsetting were performed using Seurat v5 (Hao et al., 2024). Starting from the total merged cell population, we first enriched for neural and glial progenitor populations by retaining cells expressing *Vimentin* (Figures S3C-G), a well-established marker of progenitor cell states (Barry and McDermott, 2005; Chen et al., 2018). From the *Vim*+ population, we performed a second round of subsetting to refine the dataset. Specifically, we retained (i) cells belonging to cluster 1 from the initial clustering analysis, which was characterized by concentrated expression of canonical NSPC markers, and (ii) cells expressing the pro-neurogenic transcription factor *Ascl1* (Figures S3H-L). In the final iteration of subsetting, cells were grouped into transcriptionally distinct clusters using unsupervised graph-based clustering applied to the top 30 principal components derived from normalized gene expression data, as indicated by the elbow plot (Figure S3M). Using a resolution of 0.2, we identified six unique clusters, which were visualized using uniform manifold approximation and projection (UMAP). Cell sex was assigned based on sex-linked gene expression, with cells expressing *Xist* alone classified as female and those expressing *Ddx3y*, *Uty* or *Eif2s3y* classified as male. A total of 4,485 cells collected from E15.5 control and prenatal maternal cold stress male and female embryos (control male n=3, female n=3; cold stress male n=4, female n=2) were utilized for downstream analysis (Figures S3N-P). Sequencing metrics for the final filtered cell population are summarized in Figure S3Q. The resulting six clusters were annotated and distinguished using established and differentially expressed marker genes uniquely enriched in each cluster (Figure 4A), which were identified using publicly available datasets (Kim et al., 2020, 2025).

**Figure 4.**
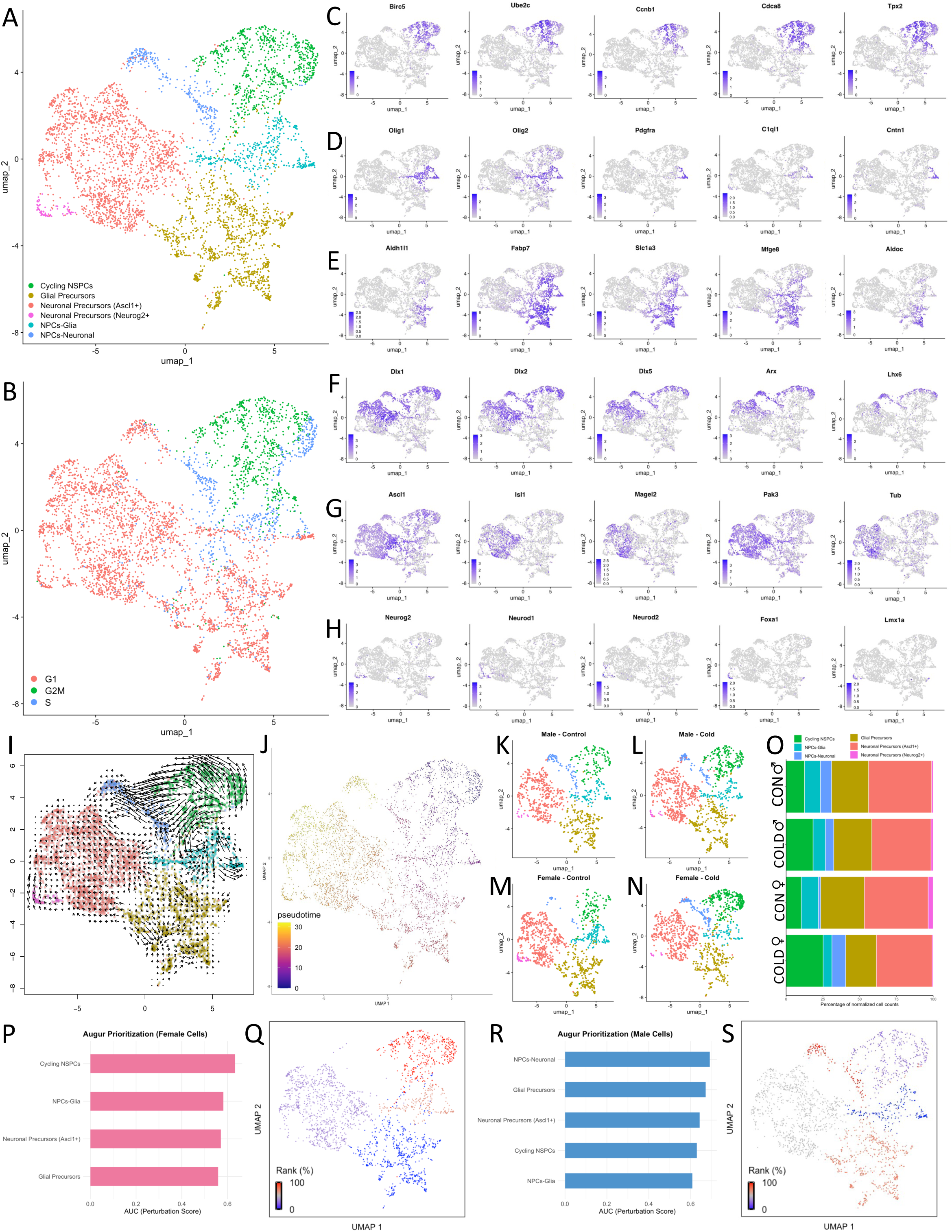
Single-cell RNA sequencing of the E15.5 hypothalamus identifies distinct subpopulations of NSPCs, where female cycling NSPCs are most responsive to prenatal maternal cold stress exposure. (A) UMAP plot of 4,485 cells collected from E15.5 control and cold stress male and female embryos (control male n=3, female n=3; cold stress male n=4, female n=2) highlight six subpopulations: cycling NSPCs, NPCs-glia, NPCs-neuronal, glial precursors, neuronal precursors (*Ascl1+*), and neuronal precursors (*Neurog2+*). (B) UMAP of cells showing cell-cycle states (G1, S, and G2/M). (C) Unique gene expression signatures of cycling NSPCs cluster. (D) Unique gene expression signatures of NPCs-glia cluster. (E) Unique gene expression signatures of glial precursors cluster. (F) Unique gene expression signatures of NPCs-neuronal cluster. (G) Unique gene expression signatures of neuronal precursors (*Ascl1+*) cluster. (H) Unique gene expression signatures of neuronal precursors (*Neurog2+*) cluster. (I) UMAP plot showing RNA velocity-predicted trajectories. (J) UMAP plot showing Pseudotime-predicted states. (K-N) Individual UMAPs showing the distribution of the cell clusters across each sex and treatment group. (O) Bar plot illustrating the relative proportion of cells in each cluster across sex and treatment group. (P) Bar plot of Augur cell-type prioritization scores for E15.5 cold stress female NSPC clusters compared to control female NSPC clusters, represented as area under the receiver operating characteristic curve (AUC) values. (Q) UMAP visualization of Augur cell-type prioritization scores ranked relative to one another for cold stress female NSPC clusters compared to control female NSPC clusters. (R) Bar plot of Augur cell-type prioritization scores for E15.5 cold stress male NSPC clusters compared to control male NSPC clusters, represented as AUC values. (S) UMAP visualization of Augur cell-type prioritization scores ranked relative to one another for cold stress male NSPC clusters compared to control male NSPC clusters.

As a first step in cluster characterization, cell cycle scoring was performed using Seurat v5 (Hao et al., 2024). Cell cycle phase assignments were determined based on the relative expression of canonical S and G2/M phase marker genes. This analysis revealed a cluster of cells in G2/M phase, indicating an actively dividing population (Figure 4B). Cells immediately diverging from this cluster were predominantly assigned to the S phase, consistent with ongoing DNA synthesis in preparation for cell division (Figure 4B). Further along the trajectory, an increasing proportion of cells were classified as G1 phase, indicating reduced proliferative activity (Figure 4B).

Consistent with its proliferative state, the G2/M-enriched cluster expressed high levels of genes associated with cell cycle progression and progenitor identity, including *Birc5*, *Ube2C*, *Ccnb1*, *Cdca8*, and *Tpx2* (Figure 4C). Based on these characteristics, this cluster was annotated as cycling NSPCs (756 cells), representing the most progenitor-like population within the dataset. Examination of cluster organization and lineage-specific marker expression revealed two transcriptionally distinct groups emerging from the cycling NSPC population, consistent with glial and neuronal lineages.

Within the glial lineage, the cluster immediately adjacent to cycling NSPCs contained cells within G2/M, S, and G1 and expressed early glial-associated markers such as *Olig1*, *Olig2*, *Pdgfr*α, *C1ql1*, and *Cntn1* (Figure 4D). This population was annotated as neural progenitor cells (NPCs)-glia (408 cells) and was interpreted as an intermediate transitional state, as it exhibited residual proliferative characteristics while concurrently upregulating early glial lineage markers. Progressing along this lineage, a subsequent cluster exhibited a predominance of G1-phase cells and expressed more mature glial precursor markers, including *Aldh1l1*, *Fabp7*, *Slc1a3*, *Mfge8*, and *Aldoc* (Figure 4E). Based on its reduced proliferative profile and enrichment of established glial precursor gene signatures, this population was annotated as glial precursors (1,137 cells).

A similar organizational pattern was observed along the neuronal lineage, whereby an intermediate cluster diverging from cycling NSPCs contained cells within G2/M, S, and G1 and expressed early neuronal markers, including *Dlx1*, *Dlx2*, *Dlx5*, *Arx*, and *Lhx6* (Figure 4F). This population was annotated as NPCs-neuronal (268 cells), consistent with its residual proliferative profile and initiation of neuronal lineage-associated transcriptional programs. Progressing along this trajectory, two transcriptionally distinct neuronal precursor populations were identified, both predominantly enriched in G1-phase cells, suggesting reduced proliferative activity and increased lineage commitment. One population expressed markers such as *Ascl1*, *Isl1*, *Magel2*, *Pak3*, and *Tub* (Figure 4G) and was annotated as neuronal precursors (*Ascl1*+) (1,839 cells), reflecting activation of gene programs associated with neuronal differentiation. The second population expressed *Neurog2*, *Neurod1*, *Neurod2*, *Foxa1*, and *Lmx1a* (Figure 4H) and was annotated as neuronal precursors (*Neurog2*+) (77 cells), consistent with a distinct neuronal differentiation trajectory. The presence of these distinct glial and neuronal lineages is consistent with previously described hypothalamic developmental trajectories (Kim et al., 2020, 2025; Fong et al., 2023).

Sequencing quality control metrics were evaluated to assess potential effects of prenatal maternal cold stress on sequencing performance and to identify possible batch effects. Unique molecular identifier (UMI) counts and the number of detected genes per cell were comparable between the two treatment groups, consistent with similar sequencing depth and library complexity (Figures S3R-S). In addition, scatter plots of UMI counts per cell versus the percentage of mitochondrial gene expression, as well as UMI counts per cell versus the number of detected genes per cell, showed substantial overlap across conditions, indicating no obvious treatment-associated differences in sequencing quality (Figures S3T-U). Together, these findings indicate that the observed transcriptional differences between conditions are unlikely to be attributable to variation in sequencing depth or library complexity. UMI counts and the number of detected genes per cell were further examined across the six identified clusters (Figures S3V-W). Since cluster sizes varied, variability in UMI and gene counts is expected, particularly in smaller clusters, due to intrinsic biological heterogeneity and the disproportionate influence of individual cells on cluster-level distributions. Importantly, as all differential analyses were conducted within clusters rather than between clusters, this variability does not impact the validity of our comparisons. Notably, mitochondrial gene expression was significantly enriched in glial precursor cells from cold stressed embryos, while NPC-neuronal cells exhibited higher mitochondrial gene enrichment under control conditions (Figure S3X). Given that mitochondrial gene expression is often associated with cellular health, we assessed a KEGG-curated apoptosis gene set (Sharma and Sampath, 2019; Kanehisa et al., 2025). Enrichment of apoptosis-related genes was observed exclusively in the neuronal precursor (*Ascl1*+) cluster under control conditions when compared with cold stress, indicating that prenatal maternal cold stress exposure does not significantly compromise NSPC viability (Figure S3Y).

Although cell cycle state and marker expression strongly suggest a progression from cycling NSPCs toward differentiated neuronal and glial lineages, these features alone cannot establish the directionality of cell state transitions. We therefore applied RNA velocity analysis, which models transcriptional dynamics based on the relative abundance of unspliced (precursor) and spliced (mature) mRNA transcripts. RNA velocity-predicted trajectories identified the cycling NSPCs cluster as the origin of NSPC differentiation, with velocity vectors diverging outward from this cluster toward the intermediate and then more mature neuronal and glial states (Figure 4I). Consistent with these predictions, pseudotime analysis (Trapnell et al., 2014) revealed that early-stage cells were enriched within the cycling NSPCs cluster, with pseudotime values progressively increasing along the inferred trajectories as cells transitioned into lineage-committed and more differentiated states (Figure 4J). Notably, neuronal precursor populations occupied the latest pseudotime state—indicating these clusters are the most developed cells at this particular developmental time-point, aligning with established hypothalamic developmental patterns in which neurogenesis precedes gliogenesis (Fong et al., 2023).

Having established the developmental trajectories and differentiation hierarchy of NSPCs, we next examined whether prenatal maternal cold stress exposure altered the distribution of cells across these transcriptional states. Cells from both male and female embryos, under control and prenatal maternal cold stress conditions, were broadly represented across all clusters; however, there were slight differences in the proportion of cells within individual clusters in both males and females (Figures 4K-O). It should be noted that these qualitative differences may also be influenced by sampling variability inherent to single-cell sequencing, as only a subset of the total cell suspension is captured and analyzed. Moreover, it is also worth noting that it can be challenging to draw robust conclusions from clusters comprised of small cell populations, such as those represented by the neuronal precursors (*Neurog2*+) cluster. Collectively, these analyses indicate that cycling NSPCs represent the most undifferentiated or stem cell-like population, giving rise to increasingly differentiated neuronal and glial lineages.

### Prenatal maternal cold stress exposure alters the transcriptomic profile of NSPCs uniquely in each sex

Interestingly, Augur cell type prioritization (Skinnider et al., 2021) revealed that female cycling NSPCs were the cells most responsive to prenatal maternal cold stress exposure (Figures 4P-Q), an effect that was not observed in males (Figures 4R-S). Given that cycling NSPCs represent the most undifferentiated or stem cell-like population that give rise to differentiated lineages and were prioritized by Augur as the most responsive cell type in females exposed to prenatal maternal cold stress, subsequent analyses focused on this cluster. To further characterize this cell population and determine how prenatal maternal cold stress impacts its transcriptional state, we analyzed differentially expressed genes (DEGs) in cycling NSPCs. We first examined basal sex differences by comparing control male and female cells (Figure 5A). This analysis identified 123 significantly upregulated and 2 significantly downregulated genes in males relative to females (Table S4). Next, we assessed the impact of prenatal maternal cold stress exposure on gene expression in male cycling NSPCs (Figure 5B), which revealed 20 significantly upregulated and 19 significantly downregulated genes in cold stress males relative to control males (Table S4). In contrast, prenatal maternal cold stress exposure drove more pronounced transcriptional responses in female cycling NSPCs (Figure 5C), with 109 significantly upregulated and 152 significantly downregulated genes in cold stress females relative to control females (Table S4). Interestingly, upregulated DEGs identified in prenatal maternal cold stress female cycling NSPCs showed substantial overlap with those identified in control males when each was compared to control females. Indeed, 51 genes were shared between control males and prenatal maternal cold stress female cycling NSPCs relative to control females. Among these were key genes such as *Dlx1*, *Dlx2*, *Dlx5*, *Meis2*, and *Epha4*. The *Dlx* family and *Meis2* are well-established regulators of GABAergic neuron specification (Kim et al., 2025), while *Epha4* plays a critical role in neurite guidance and circuit assembly (Richter et al., 2007; Cramer and Miko, 2016). The shared upregulation of these genes suggests convergence on developmental programs governing neuronal specification and connectivity.

**Figure 5.**
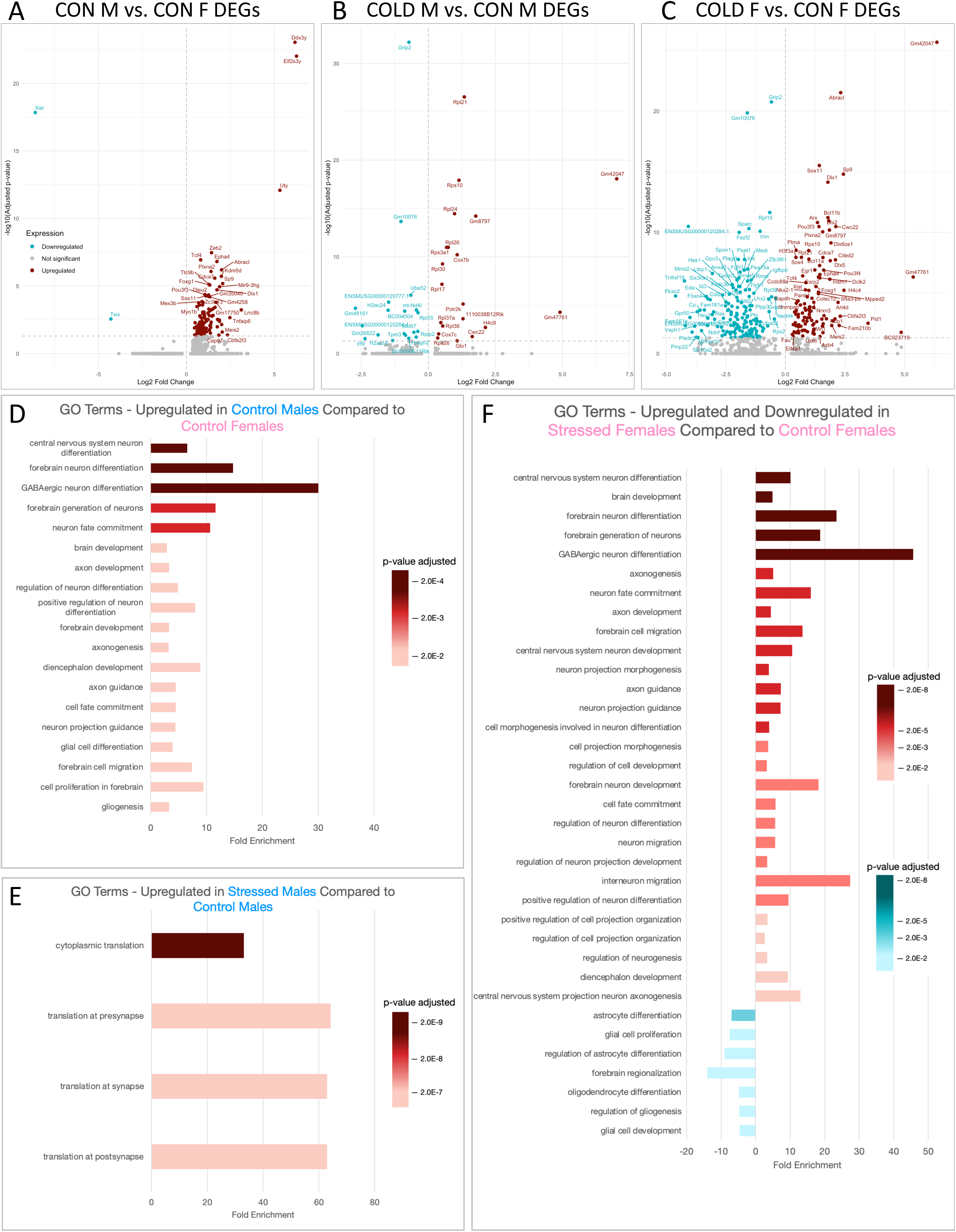
Maternal cold stress exposure alters the expression profile of female cycling NSPCs, with over-representation analysis revealing upregulation of pathways related to neuron development and GABAergic neuron differentiation. (A) Volcano plot depicting DEGs for E15.5 control male cycling NSPCs compared to control female cycling NSPCs. (B) Volcano plot depicting DEGs for E15.5 cold stress male cycling NSPCs compared to control male cycling NSPCs. (C) Volcano plot depicting DEGs for E15.5 cold stress female cycling NSPCs compared to control female cycling NSPCs. (D) GO ORA terms identified for E15.5 control male cycling NSPCs compared to control female cycling NSPCs. (E) GO ORA terms identified for E15.5 cold stress male cycling NSPCs compared to control male cycling NSPCs. (F) GO ORA terms identified for E15.5 cold stress female cycling NSPCs compared to control female cycling NSPCs.

Importantly, these transcriptional changes were validated in an independent control female NSPC replicate dataset (Figures S4A-K). In this dataset, 226 genes were significantly upregulated, and 52 genes were significantly downregulated in control male cycling NSPCs relative to control female cycling NSPCs (Figure S4L, Table S4). Similar to the prior dataset, prenatal maternal cold stress exposure also drove more pronounced transcriptional responses in female cycling NSPCs, with 281 genes significantly upregulated and 1,736 genes significantly downregulated in cold stress females relative to control females (Figure S4M, Table S4). Notably, 147 upregulated DEGs were shared between control males and prenatal maternal cold stress females relative to control females, including several of the same key regulators (e.g., *Dlx1*, *Dlx2*, *Dlx5*, and *Epha4*). Although the number of shared genes differed from the discovery dataset (i.e., 51 genes overlapped), the directionality and overall pattern of transcriptional convergence were consistent across datasets. By confirming these shared signatures in an independent cohort, we strengthen confidence that the observed transcriptional differences reflect the biology rather than technical or sampling artifacts.

We also incorporated E15 and E15.5 male and female wild-type hypothalamic cells from two published datasets, Kim et al. (2020) and Kim et al. (2025), alongside our E15.5 male and female hypothalamic datasets for further comparison of male cycling NSPCs to female cycling NSPCs (Figure S5, Table S5). In this analysis, 24 genes were significantly upregulated and 4 genes were significantly downregulated in male cycling NSPCs relative to female cycling NSPCs (Figure S5A-L, Table S5), including several of the same key regulators (e.g., *Dlx1*, *Dlx5*, *Meis2,* and *Epha4*). Again, although the number of shared genes differ from the discovery dataset, the overall pattern of transcriptional changes is consistent when assessing cycling NSPCs from Kim et al. (2020) and Kim et al. (2025) combined with cycling NSPCs datasets generated in this study, thereby helping strengthen the observed sex differences in transcriptional signatures seen between male cycling NSPCs and female cycling NSPCs.

To provide biological context for these transcriptional changes, Gene Ontology (GO) Over-Representation Analysis (ORA) was performed. Comparison of control males to control females revealed significant enrichment of pathways related to neuron development (e.g., neuron fate commitment, regulation of neuron differentiation, etc.), neuron morphology (e.g., axon development, axon guidance, etc.), and GABAergic neuron differentiation (Figure 5D, Table S4), which was also seen when wild-type cycling NSPCs from Kim et al. (2020) and Kim et al. (2025) were combined and analyzed with male and female cycling NSPCs datasets generated in this study (Figure S5M, Table S5). In contrast, comparison of prenatal maternal cold stress males to control males showed enrichment of pathways associated with translation at the pre-synapse, synapse, and post-synapse (Figure 5E, Table S4). Intriguingly, enriched pathways in prenatal maternal cold stress females closely resembled those observed in control males relative to control females, with prominent enrichment of neuron development, neuron morphology, and GABAergic neuron differentiation (Figure 5F, Table S4). Notably, the enriched GO ORA terms were influenced in part by the recurrent upregulation of *Dlx1*, *Dlx2*, *Dlx5*, *Meis2*, and *Epha4*, reinforcing that these transcriptional shifts reflect coordinated activation of neuronal specification and morphogenesis programs and further supports the concept that prenatal maternal cold stress exposure in females induces developmental programs that more closely resemble control males.

To provide spatial context and validate our scRNAseq findings *in vivo* within the developing hypothalamus, we performed fluorescent *in situ* hybridization (FISH) on serial sections spanning the rostral-caudal axis of the E15.5 hypothalamus. Using QuPath to quantitatively assess RNAscope images, where one puncta or dot equates to one individual RNA transcript, we focused our analyses on *Dlx2* and *Dlx5* expression, prominent transcription factors implicated in GABAergic neuron specification pathways that were highlighted by GO ORA. *Dlx2* and *Dlx5* were identified in our DEG analyses as being upregulated in both control males relative to control females (Figure 5A) and prenatal maternal cold stress females relative to control females (Figure 5C). Quantification of *Dlx2* and *Dlx5* expression in coronal sections across the E15.5 hypothalamus highlighted altered expression in the rostral PVN. Frequency distribution analysis showed that *Dlx2* expression was impacted by prenatal maternal cold stress in both a sex- and puncta per cell (bin)-dependent manner (Figures 6A-E, Table S6; sex x condition x bin interaction F_(20,252)_=3.095, p<0.0001). Interestingly, control females were found to have significantly more cells with no *Dlx2* puncta per cell and 1–5 *Dlx2* puncta per cell relative to control males (Figure 6E, Table S6; 0 *Dlx2* puncta per cell p<0.0001; 1–5 *Dlx2* puncta per cell p=0.0128), while control males were found to have significantly more cells with 11–15 and 16– 20 *Dlx2* puncta per cell relative to control females (Figure 6E, Table S6; 11–15 *Dlx2* puncta per cell p=0.0351; 16–20 *Dlx2* puncta per cell p=0.0305). Although comparison of control males and prenatal maternal cold stress males showed no significant changes in *Dlx2* expression (Figure 6E, Table S6), prenatal maternal cold stress females were found to have significantly fewer cells with no *Dlx2* puncta per cell and 1–5 *Dlx2* puncta per cell (Figure 6E, Table S6; 0 *Dlx2* puncta per cell p<0.0001; 1–5 *Dlx2* puncta per cell p=0.0099) and significantly more cells with 16–20 and 21–25 *Dlx2* puncta per cell relative to control females (Figure 6E, Table S6; 16–20 *Dlx2* puncta per cell p=0.0373; 21–25 *Dlx2* puncta per cell p=0.0470).

**Figure 6.**
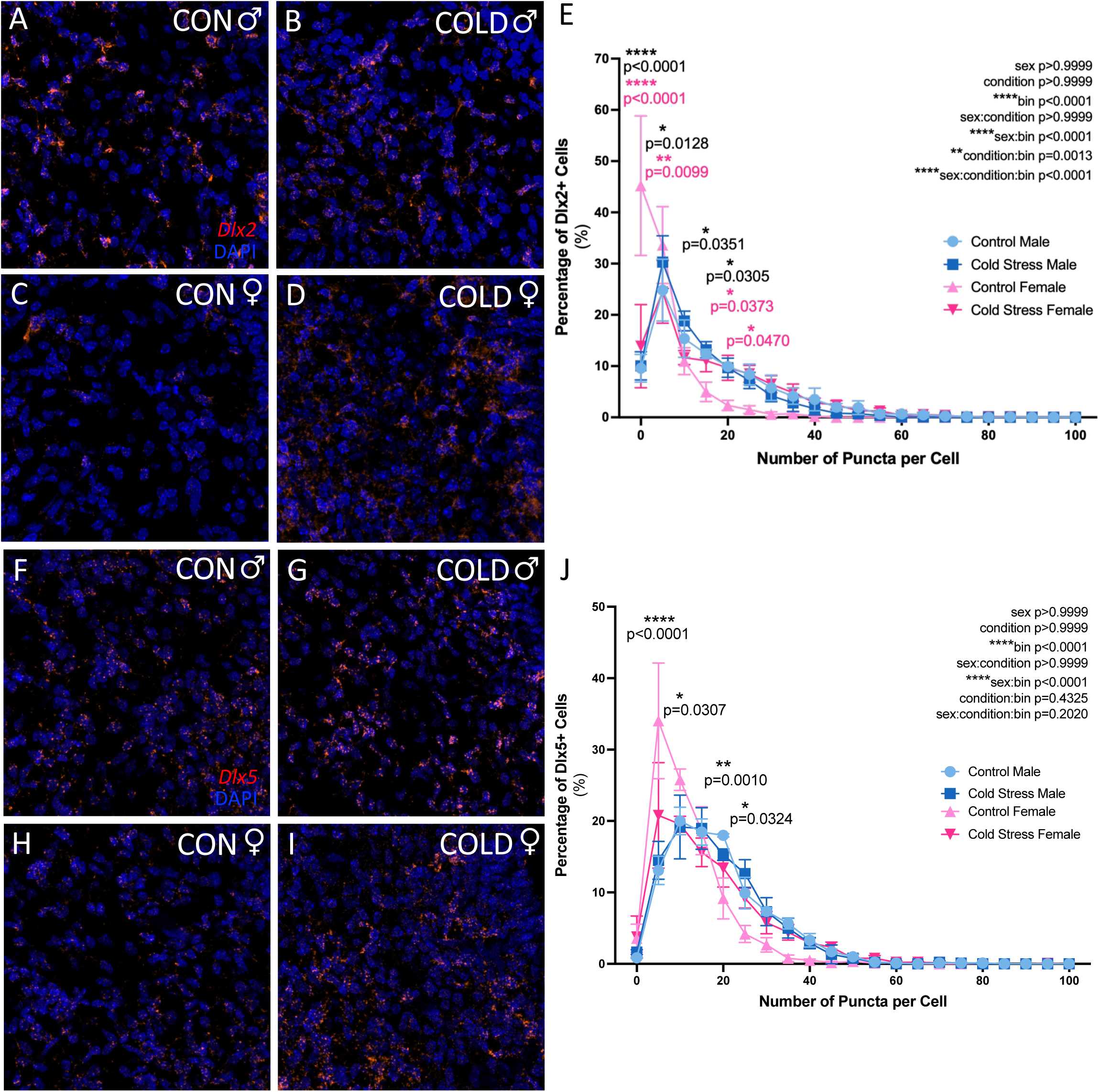
Spatial validation of DEG signatures driven by maternal cold stress exposure in the E15.5 hypothalamus. (A-D) Representative images of *Dlx2* expression and DAPI staining in the rostral PVN of the hypothalamus in male and female embryos. (E) Frequency distribution of *Dlx2* puncta per cell, comparing control and cold stress male and female embryos, displayed as the percentage of *Dlx2*+ cells binned by puncta number and normalized to the total number of cells per image. Data represent means ± SEM and were analyzed using a mixed effects model with litter incorporated as a random effect, where post hoc tests were evaluated using a Bonferroni correction (n=4 embryos, N=2-3 litters). (F-I) Representative images of *Dlx5* expression and DAPI staining in the rostral PVN of the hypothalamus in male and female embryos. (J) Frequency distribution of *Dlx5* puncta per cell, comparing control and cold stress male and female embryos, displayed as the percentage of *Dlx5*+ cells binned by puncta number and normalized to the total number of cells per image. Data represent means ± SEM and were analyzed using a mixed effects model with litter incorporated as a random effect, where post hoc tests were evaluated using a Bonferroni correction (n=3-4 embryos, N=2-3 litters). (E, J) Each dot represents a single, independent embryo (n=3-4 embryos). Dots of same color represent embryos from the same litter, while dots of different color represent embryos from different litters (N=2-3 independent litters).

Frequency distribution analysis showed that *Dlx5* expression was not impacted by prenatal maternal cold stress (Figure 6F-J, Table S6; condition F_(1,210)_<0.0001, p=1); although *Dlx5* expression differed between the sexes in a puncta per cell (bin)-dependent manner (Figure 6F-J, Table S6; sex x bin interaction F_(20,210)_=3.901, p<0.0001). Interestingly, control females were found to have significantly more cells with 1–5 and 6–10 *Dlx5* puncta per cell relative to control males (Figure 6J, Table S6; 1–5 *Dlx5* puncta per cell p<0.0001; 6–10 *Dlx5* puncta per cell p=0.0307), while control males were found to have significantly more cells with 16–20 and 21– 25 *Dlx5* puncta per cell relative to control females (Figure 6J, Table S6; 16–20 *Dlx5* puncta per cell p=0.0010; 21–25 *Dlx5* puncta per cell p=0.0324).

Taken together, DEG analysis, GO ORA, and spatial evaluation *in vivo* within the developing hypothalamus using FISH suggest that prenatal maternal cold stress exposure elicits unique transcriptional responses in cycling NSPCs in each sex, with females showing a greater magnitude of transcriptional change and altering pathways involved in neuron development, neuron morphology, and GABAergic differentiation.

### Exposure to prenatal maternal cold stress drives persistent changes in expression profiles in female GABAergic neurons

Given that our cycling NSPC analyses implicated pathways associated with GABAergic neuron differentiation, we wondered how these changes in the transcriptional profile of cycling NSPCs, alongside prenatal maternal cold stress exposure itself, would impact GABAergic neurons in the developing hypothalamus. To assess GABAergic neurons, we selected cells expressing *Gad1* and/or *Gad2*, which encode glutamate decarboxylase (GAD)—the enzyme responsible for GABA synthesis (Erlander et al., 1991). UMAP clustering of these cells revealed five transcriptionally distinct groups (Figure 7A), which were seen across both sexes and treatment conditions (Figures 7B-E). As expected, *Gad2* expression was broadly detected across the GABAergic dataset (Figure 7F). Despite this shared identity, each cluster exhibited distinct marker gene expression profiles, with cluster 0 enriched for *Hmx2* (Figure 7G), cluster 1 for *Zic1* (Figure 7H), cluster 2 for *Lhx6* (Figure 7I), cluster 3 for *Top2a* (Figure 7J), and cluster 4 for *Meis2* (Figure 7K), indicating molecular heterogeneity within developing hypothalamic GABAergic neurons.

**Figure 7.**
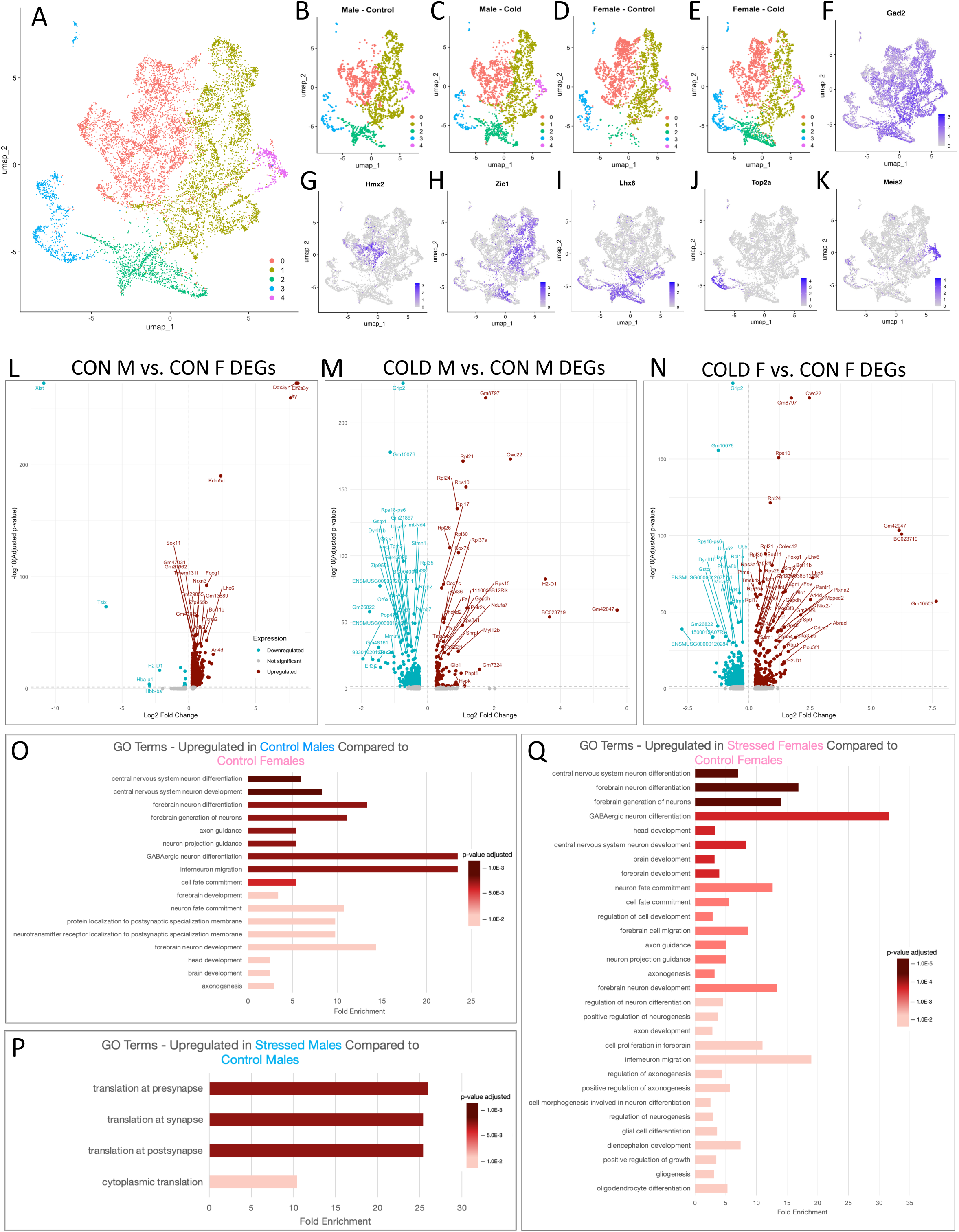
Maternal cold stress exposure alters the expression profile of female GABAergic neurons, with over-representation analysis revealing upregulation of pathways related to neuron development and GABAergic neuron differentiation. (A) UMAP plot of 9,660 *Gad1+* and/or *Gad2+* GABAergic cells collected from E15.5 control and cold stress male and female embryos highlight five subpopulations. (B-E) Individual UMAPs showing the distribution of the cell clusters across each sex and treatment group. (F) UMAP highlighting expression of *Gad2*. (G-K) Unique gene expression signatures for each of the five clusters. (L) Volcano plot depicting DEGs for E15.5 control male GABAergic neurons compared to control female GABAergic neurons. (M) Volcano plot depicting DEGs for E15.5 cold stress male GABAergic neurons compared to control male GABAergic neurons. (N) Volcano plot depicting DEGs for E15.5 cold stress female GABAergic neurons compared to control female GABAergic neurons. (O) GO ORA terms identified for E15.5 control male GABAergic neurons compared to control female GABAergic neurons. (P) GO ORA terms identified for E15.5 cold stress male GABAergic neurons compared to control male GABAergic neurons. (Q) GO ORA terms identified for E15.5 cold stress female GABAergic neurons compared to control female GABAergic neurons.

Similar to our cycling NSPCs, we first examined basal sex differences by comparing control male and female GABAergic neurons (Figure 7L). This analysis identified 919 significantly upregulated and 26 significantly downregulated genes in control males relative to females (Table S7). Next, we assessed the impact of prenatal maternal cold stress on gene expression in male GABAergic neurons (Figure 7M), which revealed 88 significantly upregulated and 610 significantly downregulated genes in cold stress males relative to control males (Table S7). In contrast, prenatal maternal cold stress drove more pronounced transcriptional responses in female GABAergic neurons (Figure 7N), with 237 significantly upregulated and 509 significantly downregulated genes in cold stress females relative to control females (Table S7).

Interestingly, upregulated DEGs identified in prenatal maternal cold stress female GABAergic neurons showed substantial overlap with those identified in control males when each was compared to control females. Indeed, 100 genes were shared between control male and prenatal maternal cold stress female GABAergic neurons relative to control female GABAergic neurons. Notably, this overlap included key developmental regulators such as *Dlx2*, *Dlx5*, and *Epha4*.

These genes are well-established drivers of GABAergic lineage specification, neuronal differentiation, and neurite guidance (Richter et al., 2007; Kim et al., 2025), and importantly, they were also among the shared regulators identified in our cycling NSPC population (Figure 5).

To provide biological context for these transcriptional changes, GO ORA was performed. Comparison of control male and female GABAergic neurons revealed significant enrichment of pathways related to neuron development (e.g., neuron fate commitment, etc.), neuron morphology (e.g., neuron projection, axon guidance, etc.), and GABAergic neuron differentiation (Figure 7O, Table S7). In contrast, comparison of prenatal maternal cold stress male and control male GABAergic neurons showed enrichment of pathways associated with translation at the pre-synapse, synapse, post-synapse, and within the cytoplasm (Figure 7P, Table S7). Interestingly, enriched pathways in prenatal maternal cold stress female GABAergic neurons closely resembled those observed in control males relative to control females, with prominent enrichment of neuronal development, neuronal morphology, and GABAergic neuron differentiation (Figure 7Q, Table S7). Although several enriched GO terms were influenced by the recurrent upregulation of *Dlx2* and *Epha4*, the contribution of *Dlx*-family regulators (e.g., *Dlx1/5*) was less prominent in the GABAergic neuron GO ORA terms compared with the cycling NSPC population. Instead, genes positioned downstream of the *Dlx* transcriptional network, including *Arx* and *Lhx6*, showed sustained representation within pathways associated with GABAergic specification and neuronal maturation. Given their established role as downstream effectors of *Dlx*-mediated developmental programs (Petryniak et al., 2007; Colasante et al., 2008), their continued involvement at this stage is consistent with the more differentiated state of this cell population. Together, these findings suggest that transcriptional programs initiated at the progenitor stage are retained during neuronal maturation but increasingly reflect the activity of downstream regulators associated with later stages of GABAergic differentiation. Consistent with patterns observed in cycling NSPCs, these results indicate that prenatal maternal cold stress exposure induces transcriptional alterations early in development that are maintained as cells mature into GABAergic neurons, suggesting that early stress-driven perturbations may leave a lasting molecular imprint on differentiated hypothalamic neurons.

### Prenatal maternal cold stress exposure alters cell-cell interactions within cycling NSPCs

Considering that prenatal maternal cold stress exposure alters the transcriptomic profile and behavior of hypothalamic NSPCs, potentially through microglia-dependent or direct mechanisms on the NSPCs themselves, we next assessed cell-cell communication using transcriptional ligand-receptor analysis (Jin et al., 2025). This analysis revealed pronounced alterations in predicted cycling NSPC communication in response to prenatal maternal cold stress exposure. In males, prenatal maternal cold stress led to a marked reduction in predicted autocrine signaling within cycling NSPCs across several key neurodevelopmental pathways, including semaphorin-plexin, neurexin, GABA, and ephrin-Eph signaling (Figure 8A). These pathways are well-established regulators of neuronal development. More specifically, semaphorin-plexin signaling plays a central role in regulating neurite guidance, dendritic arborization, and synapse formation (Kolodkin et al., 1993; Alto and Terman, 2017). Ephrin-Eph interactions enable contact-dependent signaling essential for neurite pathfinding and neuronal migration (Richter et al., 2007; Cramer and Miko, 2016). Neurexin signaling promotes synapse formation and postsynaptic differentiation, particularly at GABAergic synapses (Graf et al., 2004; Südhof, 2017), while GABA signaling contributes to early neuronal network formation (Kasyanov et al., 2004; Chalifoux and Carter, 2011). The coordinated reduction in these pathways indicates attenuation of intrinsic signaling programs within male cycling NSPCs under prenatal maternal cold stress conditions. In contrast, prenatal maternal cold stress female cycling NSPCs exhibited increased autocrine signaling through these same pathways relative to control females, including semaphorin-plexin, neurexin, GABA, and ephrin-Eph signaling (Figure 8A). This pattern suggests an enhancement of intrinsic communication networks governing neuronal guidance and synaptic organization within female cycling NSPCs following prenatal maternal cold stress exposure. These findings demonstrate that prenatal maternal cold stress exposure impacts cycling NSPC communication networks uniquely in each sex, with prominent changes seen between cycling NSPC-cycling NSPC signaling and occurring in opposite directions between males and females.

**Figure 8.**
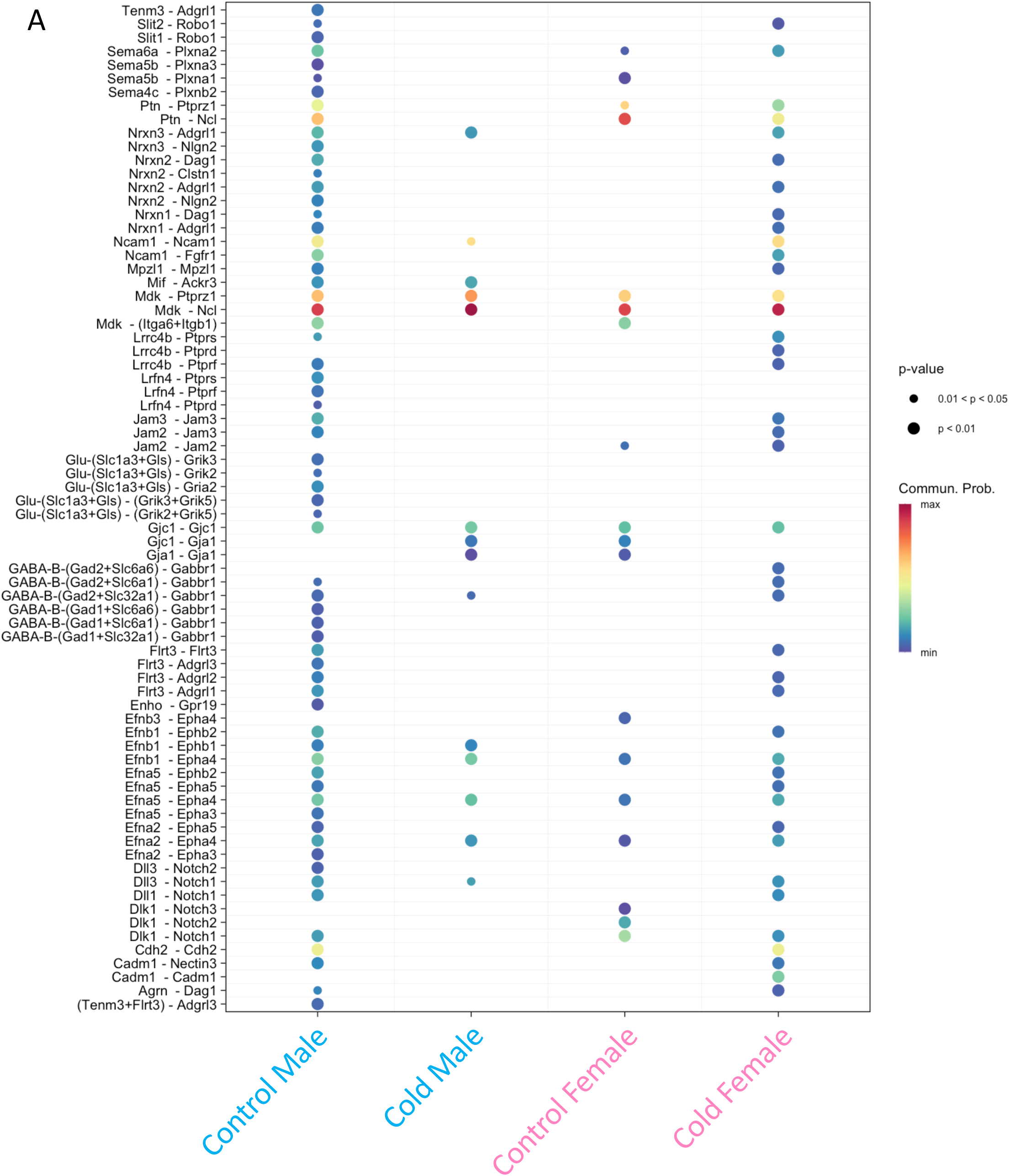
Cell-cell interaction analysis suggests that maternal cold stress exposure predominantly affects cycling NSPC-cycling NSPC signaling, with male and female embryos exhibiting changes in opposite directions. (A) Cycling NSPCs (ligand) to cycling NSPCs (receptor) signaling in male and female control and prenatal maternal cold stress conditions. Each dot represents a specific ligand-receptor interaction (y-axis). Dot size reflects the statistical significance of the interaction, with larger dots indicating more significant interactions. Dot color represents the predicted communication probability.

## DISCUSSION

In this study we demonstrated that prenatal maternal cold stress exposure does not globally impair NSPC proliferation, stem-like potential, or broad glial lineage allocation in the developing hypothalamus, indicating preserved core progenitor competence. Instead, it induces unique adaptations of NSPC transcriptional programs and developmental trajectories that differ between the sexes. Specifically, cycling NSPCs displayed baseline transcriptional sex differences that were further accentuated by prenatal maternal stress exposure. In males, maternal stress was associated with reduced autocrine NSPC signaling (scRNAseq) and altered neuronal output (*in vitro* differentiation assay). In females, maternal stress enhanced autocrine signaling, promoted GABAergic differentiation, and increased dendritic complexity, consistent with accelerated maturation. Together, these findings suggest that prenatal maternal stress recalibrates developmental trajectories within the hypothalamic NSPC niche uniquely in each sex.

Maternal stress is challenging to model in animals as pregnant women can be exposed to a range of physiological (e.g., infection) and psychological (e.g., emotional stress) stressors. A common feature reported across various maternal stress animal models is elevated maternal CORT and altered immune signaling. For example, maternal immune activation (MIA; induced using poly I:C or LPS) and restraint stress have both been shown to elevate maternal CORT and circulating cytokine levels in rodents (Cui et al., 2011; Liu et al., 2014; Delorme et al., 2021; Tang et al, 2022). This highlights commonalities between physiological (MIA) and psychological (restraint) maternal stress paradigms. Although cold exposure has been used alongside other psychological stressors such as restraint in maternal chronic unpredictable stress (CUS) models (Weinstock et al., 2017), maternal cold stress also shows similarities to MIA. Indeed, prenatal maternal cold exposure results in increased maternal CORT, placental inflammation, and altered cytokine and chemokine signaling in the embryonic brain (Lian et al., 2017; Rosin et al., 2021). Prenatal maternal cold stress exposure also drives long-term disruptions in social behaviors in both male and female offspring, although this differs between sexes, where males show microglial-dependent mechanisms and females appear to develop social deficits regardless of whether microglia are present or absent (Rosin et al., 2021). Despite well documented sex differences in brain organization and stress responses (Bowman et al., 2004; Ingalhalikar et al., 2014; Sutherland and Brunwasser, 2018; Parsaei et al., 2023; Simms, 2026), NSPC proliferative and stem-like capacity remained comparable across sexes in response to prenatal maternal cold stress exposure when examined using an *in vitro* neurosphere assay. This contrasts with other brain regions and models where prenatal stress variably alters proliferation, highlighting the importance of experimental context, including stressor type and regional NSPC heterogeneity (Kippin et al., 2004; Rosin et al., 2021; Wang et al., 2022). Preserved proliferative capacity suggests that hypothalamic NSPCs may be intrinsically buffered against certain environmental stressors, potentially reflecting the early developmental priority of establishing essential circuits regulating homeostasis. However, it is important to acknowledge the limitations of the neurosphere assay, as NSPCs are taken out of the embryonic brain which can impact signaling from surrounding cells and result in a loss of environmental context. Although other studies have shown similarities in NSPC proliferation and responses between the neurosphere assays and *in vivo* analyses (Nesan et al., 2021), this has not yet been validated in the context of prenatal maternal cold stress. Therefore, future experiments examining NSPC proliferation *in vivo* within the embryonic brain are required to support our findings herein.

Consistent with this resilience, no changes were observed in OPC, oligodendrocyte or astrocyte abundance, indicating stable glial lineage allocation. Importantly, preserved glial abundance does not preclude functional alterations. Glial cells play critical roles in supporting neuronal development and circuit function (Bradl and Lassmann, 2010; Murphy-Royal et al., 2015; Boddum et al., 2016; Masumura et al., 2025; Wei and Morrison, 2025), and prenatal stress has been shown to disrupt these functions, including impairing myelin production (Lu et al., 2022; Crombie et al., 2023). Thus, while gross glial composition appears to be preserved in response to prenatal maternal cold stress exposure, subtle functional changes cannot be excluded. Together, these findings suggest that hypothalamic progenitor maintenance and glial lineage allocation appear robust, potentially reflecting region-specific mechanisms that preserve developmental stability during environmental challenge.

Despite this apparent developmental stability, sex remains a major intrinsic biological variable that can also influence brain function and organization (Ingalhalikar et al., 2014; Parsaei et al., 2023; Simms, 2026). As the first surge in testosterone occurs starting on day 17 of gestation in rats (Ward et al., 2003), which is equivalent to approximately E15.5 to E16 in mice, exposure to sex hormones could play a role in cellular responsiveness to maternal stress and contribute to sex differences in E15.5 male and female embryos. Indeed, we observed distinct changes in each sex, indicating that male and female hypothalamic neural cells may differentially respond to prenatal stress. In males, prenatal maternal cold stress exposure resulted in an increase in TUJ1+ neurons derived from E15.5 NSPCs. Interestingly, prior studies on the prenatal maternal cold stress model demonstrated that the same paradigm reduces the number of oxytocin neurons within the E15.5 PVN (Rosin et al., 2021). Since oxytocin neurons are generated during a defined developmental window that precedes E15.5 (Grinevich et al., 2015), their reduction at E15.5 suggests that early lineage-specific neurogenesis may be disrupted or delayed in response to prenatal maternal cold stress exposure. When considered alongside the increased number of TUJ1+ neurons observed in our study, this pattern is consistent with a temporally shifted neurogenic trajectory. Specifically, an early stress-induced deficit in oxytocin neuron production may result in compensatory or delayed neuronal output later in embryogenesis, which could lead to an accumulation of immature TUJ1+ neurons. Given that TUJ1 labels early post-mitotic neurons rather than fully mature neuronal subtypes, the increase in TUJ1+ cells observed in males may reflect an accumulation of immature neurons that have exited the cell cycle but have not yet acquired terminal subtype identity (Zhang and Jiao, 2015). Therefore, the elevated TUJ1 signal may represent a redistribution in the timing and composition of neuronal production within the male hypothalamus to offset earlier deficits; however, this is speculative and would need to be examined *in vivo* within the embryonic brain across a range of time-points using birth-dating experiments and immunofluorescence staining.

In female embryos, prenatal maternal cold stress exposure induced a shift in cycling NSPC fate and neuronal morphology. Transcriptional analyses supported this, showing enrichment of pathways related to GABAergic differentiation and neuronal morphogenesis. These molecular changes were reflected morphologically, with *in vitro* analysis revealing enhanced dendritic arborization, indicative of more mature neurons (Tronel et al., 2010; Dicke and Roth, 2016; Forrest et al., 2018). Notably, the phenotype of prenatal maternal cold stress exposed female embryos qualitatively resembled that of control males. This is consistent with evidence that males display higher baseline hypothalamic GABAergic activity (Grattan and Selmanoff, 1997; Searles et al., 2000). Importantly, while such sex differences in GABAergic signaling are well established in adult systems (Grattan and Selmanoff, 1997; Searles et al., 2000), our findings suggest that these divergences may emerge during fetal neurodevelopment and can be modulated by prenatal environmental conditions. However, as these *in vitro* assays and transcriptomic analyses are correlative in nature, future experimentation targeted at examining the timing of neuronal differentiation, specification towards GABAergic lineages, and morphological analysis of neuronal dendrites *in vivo* within the embryonic brain are required to support these findings. Moreover, the inclusion of additional time-points evaluating the impact of the prenatal maternal cold stress paradigm on neurodevelopment are necessary to verify whether the sex differences observed herein truly exist or if the same alterations can be observed in each sex but at earlier or later developmental ages.

These early cellular and structural changes may have downstream functional consequences. In this context, the enhanced GABAergic differentiation and neuronal maturation observed in female embryos exposed to prenatal maternal cold stress may provide an early cellular basis for a shift toward more male-like behavioral patterns. Indeed, female offspring exposed to prenatal maternal cold stress show decreased resting time in the open field, displaying activity levels more similar to control males, while this same behavioural change was not observed in prenatal maternal cold stress exposed male offspring (Rosin et al., 2021). More broadly, stress has been shown to produce region-specific structural remodeling of neuronal architecture, often accompanied by behavioral changes (Vyas et al., 2002; Brunson et al., 2005; Liston et al., 2006; McEwen et al., 2016). However, these effects have typically been described following neonatal or adult stress exposure, whereas our findings demonstrate that structural remodeling may already be detectable at E15.5, immediately following prenatal maternal cold stress exposure.

This suggests that neuronal architecture may be sensitive to environmental perturbation earlier than previously examined, and that stress-induced structural changes could begin during the initial stages of circuit formation, leading to long-term alterations in neurodevelopment.

However, these points remain hypothetical and would need to be tested *in vivo* by examining neuronal connectivity using brain clearing or other experimental assays which can examine structural remodeling. Similarly, further assessment of behavior in prenatal maternal stress exposed offspring, including various aspects of social interaction and perhaps sex behaviors, would provide context for the cellular changes observed herein.

It is important to note, however, that the response of female cycling NSPCs to prenatal maternal cold stress exposure cannot be fully explained as a simple shift toward a more “male-like” state. A substantial number of genes were uniquely downregulated in prenatal maternal cold stress females—changes not observed in control males when compared to control females, suggesting that this response also involves distinct molecular adaptations. Our data further characterized baseline transcriptional programs in control male and female embryonic hypothalamic cycling NSPCs at E15.5, revealing 123 genes significantly upregulated and 2 genes downregulated in males relative to females. By explicitly accounting for embryo sex in our own study, and through the segregation and re-analysis of male and female hypothalamic cells independently using published datasets (Kim et al., 2020, 2025), our study extends prior work by demonstrating that hypothalamic progenitors may exhibit intrinsic molecular differences between males and females under homeostatic conditions. These intrinsic mechanisms that generate early baseline sex differences may establish the transcriptional architecture of cycling NSPCs and influence their sensitivity to environmental perturbations. While intrinsic variations in the genetic and/or epigenetic makeup of male and female embryos could be present from conception, it is important to also acknowledge that exposure to sex hormones, such as the testosterone surge around E15.5/E16 (Ward et al., 2003), may be the modifier driving these differences. Together, these findings support a model in which prenatal maternal cold stress exposure may enhance GABAergic innervation and promote structural remodeling of hypothalamic circuits in female embryos, potentially driving developmental trajectories toward what is more commonly seen in males, while simultaneously engaging unique, female-specific molecular responses. However, as the prenatal maternal cold stress scRNAseq datasets were generated using pooled embryos, which is a limitation of the current study, replication of these datasets and spatial validation *in vivo* are required to support our findings and proposed model.

Beyond cell-intrinsic changes, ligand-receptor analysis revealed unique alterations in cycling NSPC communication that differed between the sexes. In males, prenatal maternal cold stress reduced autocrine signaling, particularly in neurexin- and ephrin-associated pathways, which regulate cell-cell communication, synaptic specification, and maturation (Graf et al., 2004; Richter et al., 2007; Cramer and Miko, 2016; Südhof, 2017). Attenuation of these pathways may delay differentiation and circuit integration despite neuronal identity acquisition. In contrast, females exhibited increased autocrine signaling through these pathways, consistent with enhanced maturation, GABAergic differentiation, and dendritic complexity (Graf et al., 2004; Cramer and Miko, 2016; Südhof, 2017). Notably, neurexins have been shown to induce postsynaptic differentiation at GABAergic synapses (Graf et al., 2004), providing a potential mechanistic basis for the observed enhancement in GABAergic synapse formation and neuronal connectivity observed in our scRNAseq analysis, further highlight the importance of cell-cell signaling, communication, and environment.

These findings identify the cycling NSPC niche as a key regulator through which early environmental stress shapes fetal brain development. Importantly, this work provides some of the earliest evidence of intrinsic, transcriptionally defined sex differences in embryonic hypothalamic cycling NSPCs, highlighting that sex-specific neurodevelopmental trajectories may be established earlier than previously appreciated (Zhou et al., 2005; Garrett and Wellman, 2009; Goldstein et al., 2010). This study further provides insight into how sex-specific vulnerability to NDDs may emerge from early-life environmental perturbations. Given that maternal stress exposure during human pregnancy is strongly associated with increased risk for NDDs (Hossain and Westerlund Triche, 2007; Buss et al., 2011; Favaro et al., 2015; Bergh et al., 2018; Marečková et al., 2019; Mareckova et al., 2020; Goldstein et al., 2021; Lu et al., 2022; Yu et al., 2023), these findings help offer a biologically grounded framework linking early environmental conditions to sex-specific neurodevelopmental outcomes.

## Acknowledgments

The authors thank Felix Ma from the Rosin Laboratory for assistance with experimental training, Harmony Fong from the Kurrasch Laboratory for critical discussions, Mark Cembrowski for kindly sharing their RNAscope hybridization oven with us, the University of British Columbia (UBC) Centre for Disease Modelling for animal care, and Tara Stach, Stephen Yu, Yvonne Chung, and Bernie Zhao at the School of Biomedical Engineering Sequencing Core for generating sequencing data and providing technical support. We are grateful to Dr. Stephane Flibotte from the UBC Bioinformatics Core Facility for guidance and support related to bioinformatics and RNA velocity analyses. Kodee Bao was supported by a Canadian Institutes of Health Research (CIHR) CGS-M award. This work was supported by a CIHR Project Grant (GR028789) to J.M. Rosin. J.M. Rosin is a Michael Smith Health Research (MSHR) BC Scholar and a Tier 2 Canada Research Chair (CRC) in Immune Regulation of Developmental Programs.

## MATERIALS AND METHODS

### Experimental model

Animal work was carried out in accordance with the guidelines and regulations of the Canadian Council of Animal Care and received prior approval from the University of British Columbia’s Animal Care Committee (protocols A21-0170 and A21-0171). Adult male and female CD1 mice (strain code 022, Charles River) were used for all experiments. Timed pregnancies were used to obtain embryonic brain samples. For embryonic staging, female mice were plug-checked in the morning and those with a vaginal plug were assigned E0.5. Pregnant female mice were housed individually and continued to receive our standard chow (LabDiet PicoLab Rodent Diet 20) and water *ad libitum* during pregnancy. For embryo collection, pregnant female mice were anaesthetized with isoflurane and immediately euthanized by cervical dislocation followed by decapitation. All mouse embryos were genotyped for sex and results were reported for both sexes.

### Mouse handling and maternal cold stress paradigm

From E11.5 to E15.5, pregnant CD1 female mice were placed in a 4°C walk-in fridge for 30 minutes daily (08:00–08:30) for five consecutive days. Pregnant mice were euthanized on the last day of cold stress (E15.5) immediately following the last cold stress exposure. During the cold stress, pregnant CD1 mice remained within their home cage and had access to our standard chow (LabDiet PicoLab Rodent Diet 20) and water *ad libitum*. Lighting conditions followed the facility light cycle (lights on at 07:00 and off at 19:00) and were maintained during the stress exposure period.

### DNA extraction and genotyping

Tail tissue was collected from all mouse embryos and was incubated overnight at 56°C in extraction buffer (100 mM NaCl, 50 mM Tris, 100 mM EDTA, and 1% SDS) with proteinase K (8 units, ∼0.4 mg/mL). Genomic DNA was isolated from the digested tissue by precipitation with saturated NaCl solution (∼6 M), followed by centrifugation at 20,000 g for 10 minutes. DNA was subsequently precipitated from the supernatant using isopropanol, and pellets were collected by centrifugation at 20,000 g for 5 minutes. Pellets were washed with 70% ethanol, air-dried at 37°C, and resuspended in 200 µL of Tris-EDTA buffer (10 mM Tris, 1 mM EDTA, pH 8) for 1 hour at 68°C. For genotyping, 1.6 µL of genomic DNA was added to OneTaq® Quick-Load® 2X Master Mix with Standard Buffer (New England Biolabs), along with 1 µM of SX primers (*Sly & Xlr*) (McFarlane et al., 2013). PCR amplification was performed using an Applied Biosystems™ MiniAmp™ Thermal Cycler (Applied Biosystems) with the following cycling conditions: an initial denaturation at 95°C for 3 minutes; 35 cycles of denaturation at 95°C for 30 seconds, annealing at 57°C for 30 seconds, and extension at 72°C for 30 seconds, followed by a final extension at 72 °C for 5 minutes. PCR products were run on a 1.5% agarose gel prepared in TAE buffer (0.4 M Tris, 0.2 M acetic acid, 10 mM EDTA, pH 8) and visualized using SmartGlow Pre-Stain for nucelic acid gels (Accuris Instruments) on a Fisherbrand™ Real-Time Electrophoresis System (Fisher Scientific). Fragment sizes were determined using a GeneRuler Ready-to-Use 100 bp DNA Ladder (Thermo Fisher Scientific).

### Neurosphere assay

The neurosphere assay was performed following protocols described previously (Nesan et al., 2020; Rosin et al., 2021). In brief, E15.5 CD1 embryos, from both control and maternal cold stress-exposed dams, were dissected and placed in 37°C warmed sterile 1X phosphate buffered saline (PBS; Gibco 10010-031, Thermo Fisher Scientific) containing penicillin-streptomycin (GIBCO 15140-122, Thermo Fisher Scientific) within a petri dish. Hypothalamic ventricular tissue was micro-dissected from the E15.5 embryos while in PBS with penicillin-streptomycin and placed into labeled microfuge tubes containing 1 mL of PBS with penicillin-streptomycin. Corresponding embryo tails were removed and placed in separate microfuge tubes for later use to genotype each embryo. Under a biological safety cabinet, the micro-dissected hypothalamic ventricular tissue was mechanically dissociated by pipetting up and down 40 times with a 1 mL pipette and then centrifuged at 300 g for 10 minutes at room temperature. The PBS was decanted, and each cell pellet was resuspended in 1 mL of 37°C warmed NeuroCult Proliferation Media (STEMCELL Technologies 05702) with 20 ng/mL each of human recombinant epidermal growth factor (EGF) and basic fibroblast growth factor (bFGF) (STEMCELL Technologies), and 2 µg/mL Heparin solution (STEMCELL Technologies 07980). The cell suspension was mechanically dissociated by pipetting up and down 20 times and centrifuged at 300 g for 10 minutes at room temperature. The cell pellet was resuspended in 300 μL of media and filtered through a 35 μm strainer (Falcon 352235). Cell number and viability was assessed using the Countess II Automated Cell Counter (Thermo Fisher Scientific) with chamber slides (Invitrogen C10228, Thermo Fisher Scientific) and Trypan Blue Stain (Invitrogen T10282, Thermo Fisher Scientific). Primary neurospheres were plated at a cell density of 5,000 cells/mL in a 24-well plate (1 mL/well) and were incubated for 10 days at 37°C with 5% CO_2_, with 50% media replenishment at day 5. At day 10, primary neurospheres were imaged and quantified. Neurospheres were then either fixed in 4% paraformaldehyde (PFA) in PBS for 20 minutes, followed by three 10-minute washes in PBS for immunohistochemical analysis, or dissociated for secondary neurosphere formation or use in the differentiation assay.

For secondary neurosphere formation, primary neurospheres were dissociated under sterile conditions by mechanically scraping the bottom of each well using a 1 mL pipette tip, followed by pipetting up and down 40 times. Cells were centrifuged at 300 g for 10 minutes at room temperature and the pellet was resuspended in 150 µL warmed NeuroCult Proliferation Media (STEMCELL Technologies 05702) by pipetting up and down 20 times. Technical replicates wells were pooled (total volume 300 µL) to ensure adequate cell counts, and the suspension was passed through a 35 µm cell strainer (Falcon 352235). Cells were then counted and cultured in warmed NeuroCult Proliferation Media (STEMCELL Technologies 05702) with 20 ng/mL each of EGF and bFGF (STEMCELL Technologies), and 2 µg/mL Heparin solution (STEMCELL Technologies 07980), as used for primary neurosphere cultures. Secondary neurospheres were replated at a density 1,000 cells/mL in a 24-well plate (1 mL/well) and were incubated for 10 days at 37°C with 5% CO_2_, with 50% media replenishment at day 5. At day 10, secondary neurospheres were imaged, quantified, and fixed in PFA as described above.

### Differentiation assay

Dissociated primary neurosphere cell suspensions, generated as described above, were cultured in warmed NeuroCult Differentiation Media (STEMCELL Technologies 05704). Cells to be used in the differentiation assay were plated at a density of 5,000 cells/mL in 24-well plates containing Laminin-coated glass coverslips at the bottom of the well (Neuvitro NC0597572, Thermo Fisher Scientific). Cells were maintained at 37°C in a humidified incubator with 5% CO_2_, with 50% media replenishment at day 5. At day 10, the Laminin slips were fixed with 4% PFA for 20 minutes, followed by two 10-minute washes in PBS in preparation for immunohistochemistry.

### Immunofluorescence staining

Neurospheres and differentiation slips (Laminin-coated glass slips) were washed 3 times for 10 minutes with 1 mL PBST (1X PBS with 0.1% Triton X-100), incubated for 30 minutes in 1X PBS with 1% Triton X-100, then blocked for 1 hour using 500 μL of 5% NDS (Sigma) in PBST at room temperature. Neurospheres and differentiation slips were then exposed to 300 μL of primary antibody, including mouse anti-TUJ1 (1:500, BioLegend 801201), rabbit anti-SOX2 (1:500, Millipore AB5603), rabbit anti-OLIG2 (1:200, Millipore AB9610), goat anti-PDGFRα (1:150, R&D Systems AF1062) or rabbit anti-GFAP (1:500, Dako Z0334), diluted in PBST and 5% NDS and left to incubate overnight at 4°C. Neurospheres and Laminin slips were washed 4 times for 10 minutes with 1 mL of PBST and exposed to 300 μL of secondary antibody, including Alexa 488 donkey anti-mouse IgG (1:200, Invitrogen A21202), Alexa 488 donkey anti-goat (1:200, Invitrogen A32814), Alexa 488 donkey anti-rabbit (1:200, Invitrogen A21206) or Alexa 594 donkey anti-rabbit (1:200, Invitrogen A21207), diluted in PBST and 5% NDS and left to incubate for 2 hours at room temperature. Samples were then washed 4 times for 10 minutes with PBST, stained with Hoescht 33342 (1:1000, Invitrogen) for 5 minutes at room temperature, washed 3 times for 10 minutes with PBST, and mounted using Aqua Poly/Mount (Polysciences Inc.).

### Imaging of neurospheres

Brightfield images used for neurosphere quantifications were acquired using a ZEISS Axiovert 5 microscope at 2.5x magnification, with imaging performed across the entire well for each sample. High-resolution images of immunofluorescence-stained spheres were captured on a ZEISS LSM900 confocal microscope and were further processed using LSM+. Brightness and/or contrast of the entire image was adjusted using ZEISS ZEN 3.9 if deemed appropriate.

### Imaging of differentiation slips

Fluorescent images were captured on a ZEISS Axioplan 2 fluorescent microscope at 20x with a ZEISS Axiocam HRm camera. For differentiation slip cell quantification, five unique, non-overlapping fields of view were imaged per slip in a consistent pattern to ensure representative sampling of cellular distribution, with each glass slip representing cells collected from a single, independent embryo. For neuron morphology analysis, five representative neurons were imaged per slip, with each glass slip representing cells collected from a single, independent embryo.

### Neuron morphology analysis

Neuronal images from differentiation slips were acquired as described above. Neuronal tracings were performed using ImageJ/FIJI software with the NeuronJ plugin (Schindelin et al., 2012). Based on the 20x objective, the image scale was calibrated to 3.09694746 pixels/µm. Tracings were initiated at the junction between the soma and the dendritic projection, and all subsequent branches were traced. Neuronal axons were not included in these tracings. Quantitative measurements derived from these tracings were recorded for downstream neuron morphology analysis.

### Single-cell RNA sequencing single-cell suspension

E15.5 CD1 embryos from control and maternal cold stress-exposed dams were dissected and placed in 37°C sterile 1X PBS. Embryo tails were collected and placed in individual microfuge tubes for subsequent genotyping (control replicate 1: n=6 pooled embryos comprised of n=3 males and n=3 females; prenatal maternal cold stress replicate 1: n=6 pooled embryos comprised of n=4 males and n=2 females; control replicate 2: n=5 pooled embryos comprised of n=5 females). Hypothalamic ventricular tissue was micro-dissected from the embryos in 37°C Hanks’ Balanced Salt Solution (HBSS; Gibco 14175-095, Thermo Fisher Scientific) and then transferred to a small petri dish containing 2 mL of 37°C culture media composed of 57.4% DMEM (Gibco 11965-092, Thermo Fisher Scientific), 28.6% F-12 (Gibco 11765-054, Thermo Fisher Scientific), 5% fetal bovine serum (FBS), 5% horse serum (HS), 2% B-27 supplement (Gibco 17504-044, Thermo Fisher Scientific), 1% N2 supplement (Gibco 17502-048, Thermo Fisher Scientific), and 1% GlutaMAX (Gibco 35050-061, Thermo Fisher Scientific). Culture media was passed through a 0.1 µm strainer prior to use. Tissues from all six embryos were pooled and mechanically dissociated using a sterile razor blade. Dissociated tissue was transferred to a 15 mL centrifuge tube, and the petri dish was rinsed with 1 mL of culture media to collect any remaining tissue, which was added to the same tube. Cell suspensions were further dissociated by pipetting up and down 40 times using a 1 mL pipette and filtered through a 70 µm strainer into a new 15 mL tube containing 7 mL HBSS with 10% FBS and centrifuged at 300 g for 5 minutes at room temperature. The cell pellet was resuspended in 1 mL HBSS with 10% FBS and mixed by gentle pipetting 20 times. The cell suspension was passed through a 40 µm cell strainer, and cell concentration and viability were assessed using Trypan Blue and manual counting with a hemocytometer (VWR 15170-208). Viable and non-viable cells were quantified by visualization on a ZEISS Axiovert 25 at 10x magnification.

### Single-cell RNA sequencing library construction

Single-cell suspensions were prepared as described above and loaded onto the 10X Genomics Chromium X Controller for droplet-based single-cell encapsulation. Libraries were generated using the Chromium GEM-X Single Cell 3′ Kit v4 (10X Genomics) following the manufacturer’s standard protocol. Briefly, viable single cells were loaded at a targeted recovery of approximately 20,000 cells per sample and partitioned into nanoliter-scale Gel Bead-in-Emulsions (GEMs) using the instrument’s microfluidic channels. Following GEM incubation, emulsions were broken and barcoded cDNA was recovered and purified using silane magnetic beads according to the manufacturer’s instructions. Full-length cDNA was amplified by PCR, and quality control was performed to assess cDNA size distribution prior to library construction. Final libraries were pooled and sequenced on an Illumina NextSeq 2000 platform using paired-end sequencing, following standard 10x Genomics Chromium GEM-X Single Cell 3′ v4 sequencing recommendations. This resulted in an average sequencing depth of 24,512 mean reads per cell for ∼15,556 cells collected from E15.5 control samples and 19,632 mean reads per cell for ∼17,961 cells collected from E15.5 maternal cold stress samples, with sequencing saturation values of 39.0–40.5%. A second control run using only female cells, performed for verification purposes, yielded an average sequencing depth of 24,259 mean reads per cell for ∼23,932 cells with a sequencing saturation of 39.5%. The library preparation and sequencing described above were conducted at the School of Biomedical Engineering (SBME) Sequencing Core at the University of British Columbia.

### Bioinformatics analysis of single-cell RNA sequencing datasets

The scRNAseq data from this study have been deposited in NCBI’s Gene Expression Omnibus (GEO) (Edgar et al., 2002) and are accessible through GEO Series accession number GSE328919 (https://www.ncbi.nlm.nih.gov/geo/query/acc.cgi?acc=GSE328919). All code used in this study is publicly available at https://github.com/RosinLabUBC/scRNAseq-code-KB. Below, we briefly describe the methodologies employed for the analyses presented in this study.

Raw sequencing reads were processed using kb-python, a wrapper around kallisto / bustools implementing the methods described in (Melsted et al., 2021; Sullivan et al., 2025), to generate spliced, unspliced, and ambiguous count matrices. The reference genome used was *mus musculus* GRCm39 (mm39) with Ensembl gene annotation version 109. The processed matrices were imported into Seurat v5.2.1 (Hao et al., 2024), and Seurat objects were created with a minimum threshold of 3 cells per gene and 200 detected genes per cell. Quality control filtering was performed to remove low-quality cells, excluding cells with fewer than 1,000 detected genes or greater than 5% mitochondrial transcript content, as these likely represented damaged cells or processing artifacts. For each dataset (control and maternal cold stress), multiple assays corresponding to total RNA, ambiguous, spliced, and unspliced transcripts were added to the Seurat objects to support downstream RNA velocity analyses. Each assay was normalized independently using SCTransform, and the SCT assay was set as the default for downstream analyses. Control and maternal cold stress datasets were subsequently merged. Dimensionality reduction was performed using principal component analysis (PCA) on SCT-normalized data, and the number of informative principal components was assessed using elbow plots. Since UMAP embeddings from the merged datasets showed strong overlap, downstream analyses were performed without batch integration.

Datasets were subsetted using an iterative filtering approach. In the first iteration, cells expressing *Vimentin* (*Vim* > 0) were extracted from the merged dataset to enrich for neural and glial progenitor populations, as *Vimentin* is a well-established marker of progenitor cell states (Barry and McDermott, 2005; Chen et al., 2018). Clustering was performed using FindNeighbors and FindClusters using the first 35 principal components and a resolution of 0.1, based on parameter optimization using clustree analysis. UMAPs were used for visualization. From the *Vim+* population, cells within cluster 1, which was defined by the concentrated expression of markers characteristic of the neural stem and progenitor populations of interest, along with cells expressing the neurogenic marker *Ascl1*, were extracted and re-clustered using the first 30 principal components at a resolution of 0.2 to generate refined neural progenitor subclusters.

Cell sex was determined using sex-linked gene expression. Cells expressing *Xist* alone were classified as female, while cells expressing *Ddx3y*, *Uty* or *Eif2s3y* were classified as male. Cells expressing both male and female markers or neither were excluded from further analysis.

Treatment condition (Control or Cold) was annotated based on sample identity metadata. Clusters were annotated based on known marker gene expression.

#### Cell Cycle Scoring

Downstream analyses included cell cycle scoring which was performed using Seurat (Hao et al., 2024). This approach leverages transcriptome-wide expression of canonical cell cycle marker genes to infer cell cycle state at the single-cell level. Canonical S phase and G2/M phase marker genes curated by Seurat (cc.genes.updated.2019), originally defined in human datasets, were converted to mouse orthologs using the gprofiler2 function gorth. These mouse gene sets were used to score each cell based on relative expression of S and G2/M markers via the CellCycleScoring function on the active SCT assay. Predicted cell cycle phases were assigned to each cell and visualized on UMAP embeddings using DimPlot.

#### RNA Velocity

RNA velocity was calculated with the velocyto.R package (La Manno et al., 2018). Velocity estimation was performed with kCells = 25, deltaT = 1, and fit.quantile = 0.02 to model gene-specific steady-state transcriptional relationships. Deviations from these steady states were used to infer the direction and magnitude of transcriptional change at the single-cell level. Velocity vectors were projected onto the UMAP embedding and visualized using a correlation-based velocity field with 200 sampled cells and square-root scaling of vector lengths. Grid-based flow visualization was performed using grid.n = 40, min.grid.cell.mass = 0.5, and arrow.scale = 3.

Cell identities were colored according to Seurat cluster assignments to match the original UMAP.

#### Pseudotime

Pseudotime trajectory analysis was performed using Monocle 3 to infer dynamic cellular progression from static single-cell transcriptomic data (Trapnell et al., 2014). A previously processed and annotated Seurat object was converted into a Monocle 3 cell data set using the as.cell_data_set function. PCA and UMAP embeddings generated in Seurat were transferred to Monocle to ensure consistency between dimensionality reductions, and cluster identities were retained. Graph-based trajectories were learned using learn_graph with partitioning disabled (use_partition = FALSE) and graph pruning disabled to retain weakly connected cell states. Cells were ordered along the trajectory using order_cells, with root nodes defined based on RNA velocity-predicted trajectories, which were obtained as described previously. Pseudotime values were projected onto UMAP embeddings for visualization.

#### Augur Analysis

Cell types responsive to prenatal maternal cold stress were identified using Augur (Skinnider et al., 2021). Female and male cells were subsetted out and were analyzed separately. For both datasets, the RNA assay was set as the default assay prior to analysis. Preprocessed Seurat objects were normalized using NormalizeData, highly variable genes were identified with

FindVariableFeatures, and gene expression values were scaled across all genes using ScaleData. Data layers were joined using JoinLayers prior to downstream analysis. When multiple datasets are merged, they are initially retained as separate data layers within the object; therefore, this step is required to consolidate them into a single unified dataset for subsequent analyses. Augur analysis was performed using the calculate_auc function, with treatment condition (control vs. cold stress) provided as the perturbation label and cluster identity used to define cell types. For each cell type, Augur repeatedly subsampled cells and trained a classifier to predict treatment status from RNA expression profiles. Default Augur sampling parameters were used (n_subsamples = 50, subsample_size = 20), and analyses were parallelized using 8 computational threads (n_threads = 8). Cell types were prioritized based on AUC, where higher AUC values indicate greater transcriptional separability between conditions. Random seeds were set to ensure reproducibility. AUC scores were visualized using ranked bar plots and projected onto UMAP embeddings to display the relative perturbation responsiveness of each cell type.

#### Differential gene expression analysis

DEG analysis was performed using Seurat (Hao et al., 2024). The RNA assay was first normalized using Seurat’s NormalizeData function. Highly variable genes were then identified with FindVariableFeatures, and gene expression values were scaled across all genes using ScaleData. DEGs were subsequently identified using the FindMarkers function in Seurat, applying a minimum log fold-change threshold of 0.25 and requiring expression in at least 10% of cells within each cluster. DEG analysis was performed in a cluster-specific manner. Baseline sex differences were assessed by comparing males and females within each cluster under control conditions (both for replicate 1 and replicate 2). E15 and E15.5 wild-type hypothalamic cells from two published datasets, Kim et al. (2020) and Kim et al. (2025), were also segregated by sex (Kim et al., 2020 GEO, GSE132355: GSM3860738 Hypothalamus, embryonic E15 time point (1); Kim et al., 2025 GEO GSE284492: GSM8685159 E15.5 Wild Type scRNA-Seq Replicate 1 and GSM8685160 E15.5 Wild Type scRNA-Seq Replicate 2) and incorporated into E15.5 male and female hypothalamic datasets described in this study for a set of comparisons between male cycling NSPCs and female cycling NSPCs (i.e., all datasets were combined for comparisons). To evaluate treatment effects, comparisons between maternal cold stress and control conditions were conducted separately within male and female (i.e., COLD vs CON in males and COLD vs CON in females) datasets for each cluster (both for replicate 1 and replicate 2).

#### Gene Ontology Over-Representation Analysis

GO ORA was conducted using the clusterProfiler package (Ashburner et al., 2000; Xu et al., 2024). For GO ORA, additional post hoc filtering was applied to our DEG lists to focus on biologically meaningful and statistically robust changes. Genes were kept if they exhibited an absolute log2 fold-change greater than 0.5 and an adjusted p-value<0.05. Adjusted p-values were calculated using Bonferroni correction based on the total number of genes in the dataset, as implemented by default in Seurat. Upregulated and downregulated genes were analyzed separately. GO ORA was then performed on the filtered DEG lists to identify significantly enriched biological processes.

To define an appropriate gene universe for each analysis, Seurat objects were subsetted according to the specific comparison being performed. All analyses were conducted using the RNA assay, with data normalized using NormalizeData, variable features identified with FindVariableFeatures and scaled using ScaleData. Data layers were merged with JoinLayers prior to downstream analyses. For each cluster of interest, the Seurat object was first subsetted by cluster identity, and the gene universe was defined in a cluster-specific manner. Genes were included in the universe if they were expressed in more than 10% of cells within the cluster, based on normalized RNA expression values. For comparisons between control males and control females, the gene universe was defined using only control cells. In contrast, for condition comparisons (cold stress vs control) within each sex, the gene universe was defined using all cells of the relevant sex, including both control and maternal cold stress conditions. This approach ensured that gene universes were tailored to both the cellular context (cluster identity) and the biological comparison being performed, while avoiding exclusion of condition-specific genes.

Gene symbols from DEG lists and the gene universe were converted to Ensembl identifiers using the bitr function from the clusterProfiler package and the org.Mm.eg.db annotation database. GO ORA was then performed using the enrichGO function with the Biological Process (BP) ontology, specifying a cluster-specific gene universe to account for differences in baseline gene expression across cell types. Multiple testing correction was applied using the Benjamini-Hochberg method and GO terms with a q-value<0.05 were considered statistically significant.

GO enrichment analyses were conducted separately for upregulated and downregulated gene sets for each cluster.

#### CellChat

Cell-cell communication was inferred using CellChat v2 (Jin et al., 2025). The NSPC scRNAseq data was processed using Seurat. Analysis was performed using the RNA assay, with datasets normalized, variable features identified, and gene expression scaled as described above. NSPCs were stratified by sex and treatment condition. NSPC cluster identities were retained and used as cell-type labels. CellChat objects were constructed using this matrix. The mouse CellChat ligand-receptor database was used for all analyses. Overexpressed signaling genes and ligand-receptor interactions were identified for each cell group, and expression data were smoothed using a curated protein-protein interaction network. Communication probabilities between sender and receiver cell populations were computed and filtered to exclude interactions involving fewer than 10 cells, which is set by default. Signaling probabilities were subsequently aggregated at the pathway level to generate global intercellular communication networks.

### Fluorescent *in situ* hybridization (FISH)

Brains were dissected from E15.5 embryos, collected in ice-cold PBS, and fixed overnight at 4°C in 4% PFA. Following fixation, embryonic brains were washed 3 times in PBS and cryoprotected by equilibration in 30% sucrose prepared in PBS overnight at 4°C. Brains were then embedded in Clear Frozen Section Compound (VWR 95057-838) and cryosectioned in the coronal plane at a thickness of 10 µm on a Leica CM1950 cryostat (Nussloch, Germany). Serial sectioned tissue containing the hypothalamus was collected over 10 slides (21 sections per slide) from a single E15.5 brain and stored in -80°C. E15.5 coronal brain sections were processed using the RNAscope™ Multiplex Fluorescent Reagent Kit v2 (Advanced Cell Diagnostics 323100) according to the manufacturer’s protocol. Briefly, the tissue sections were thawed overnight at room temperature and baked for 60 minutes at 60°C using the “Bake” setting on the ACD HybEZ™ II Hybridization System (Advanced Cell Diagnostics). Sections were rehydrated in PBS and post-fixed in 4% PFA for 30 minutes at room temperature. Slides were then dehydrated through an ascending ethanol series (50%, 70%, and 100% ethanol), followed by a second incubation in fresh 100% ethanol, air drying for 5 minutes, then incubated in hydrogen peroxide for 10 minutes at room temperature. Slides were washed with distilled water before target retrieval was performed by incubating sections in boiling target retrieval buffer for 5 minutes.

Sections were subsequently washed in distilled water, dehydrated in 100% ethanol for 3 minutes, and treated with Protease III for 30 minutes at 40°C under humidified conditions using the “RNAscope” setting on the ACD HybEZ™ II Hybridization System. RNAscope probes targeting *Dlx2* (Advanced Cell Diagnostics 555951-C3) and *Dlx5* (Advanced Cell Diagnostics 478151-C3) were hybridized to the tissue for 2 hours at 40°C under humidified conditions. Following probe hybridization and subsequent washes in wash buffer, slides were stored overnight at room temperature in 5X saline-sodium citrate (SSC) buffer. The 5X SSC buffer consisted of 0.75 M NaCl (Sigma-Aldrich S5886-1KG) and 0.075 M sodium citrate tribasic dihydrate (Sigma-Aldrich C8532-500G) prepared in distilled water and adjusted to pH 7.0. The following day, tissue sections were then incubated in amplification reagents and TSA Vivid fluorophores, with washes in wash buffer between incubations. Nuclei were counterstained with DAPI for 30 seconds and slides were mounted using Aqua Poly/Mount (Polysciences Inc.).

### Imaging FISH staining

High-resolution images of E15.5 control and prenatal maternal cold stress male and female FISH-stained hypothalamic coronal sections were captured on a ZEISS LSM900 confocal microscope at 20x with 2.0x digital zoom and were further processed using LSM+. Brightness and/or contrast of the entire image was adjusted using ZEISS ZEN 3.9 if deemed appropriate.

### QuPath analysis

Acquired images were saved as .*czi* files and imported into QuPath open-source software for analysis (Bankhead et al., 2017). Image analysis was performed in accordance with protocols provided by Advanced Cell Diagnostics and guidance from the QuPath user documentation and discussion forum. Briefly, regions of interest were annotated for each image; in this study, the entire image was selected as the region of interest. Cell detection was performed using the DAPI channel, with parameters optimized to accurately delineate nuclear boundaries. These parameters included setting the *Requested pixel size* to 0 µm, *Background radius* to 0 µm, disabling *opening by reconstruction*, and setting the *Maximum area* to 200 µm^2^. Detection thresholds were visually optimized for accurate nuclear identification and then applied consistently across subsequent images with identical imaging conditions. Cells that were inaccurately detected (e.g., merged nuclei counted as a single cell) were manually excluded from further analysis by selecting and deleting the corresponding annotations. Subcellular detection was subsequently performed on each image. Parameters included enabling *Split by intensity* and *Split by shape*, with detection thresholds adjusted to optimize identification of puncta. Once optimized, these parameters were kept constant across images to ensure consistency.

### Quantification and Statistical Analysis

Comparisons between control and prenatal maternal cold stress male and female groups were performed using mixed effects model with litter incorporated as a random effect. A two-tailed unpaired t-test within each sex was only employed for secondary dendrite length, where an interaction between sex and condition was identified in the mixed effects model.

#### Neurosphere quantification

Brightfield images of neurosphere cultures were acquired as described above. Quantification was performed for each well by measuring neurosphere diameter, which was determined by manually drawing a line across the diameter of each sphere using the measurement tool in ZEN Microscopy Software (Zeiss). Only spheres with a diameter of ≥ 40 µm were included in the analysis. Quantitative results (primary neurospheres: n=8-15 embryos; secondary neurospheres: n=7-12 embryos per sex/treatment from N=3 independent dams/litters) are presented as means ± SEM. Each data point represents neurospheres derived from a single, independent embryo. For each embryo, NSPCs were plated in technical replicates, which were averaged to generate one biological data point per embryo. Statistical analyses were performed using a mixed effects model with litter incorporated as a random effect, where post hoc tests were evaluated using a Bonferroni correction in R and data was plotted using GraphPad Prism 9 (GraphPad Software). Statistical significance was defined as p<0.05.

#### Differentiation coverslip quantification

Images of differentiated NSPC cultures were acquired as described above. For each image, cell counts were performed using Adobe Photoshop to quantify TUJ1+ neurons, OLIG2+ OPCs, OLIG2+/PDGFRα+ double-positive oligodendrocytes, and GFAP+ astrocytes and normalized to total Hoechst+ cells captured in the image. For each glass coverslip, five non-overlapping fields of view were imaged and quantified, and values were averaged to generate a single data point per coverslip. NSPCs were collected from two embryos per sex/treatment/litter (n=6 total embryos per sex/treatment from N=3 independent litters), with one coverslip generated from a single, independent embryo. Quantitative results are presented as means ± SEM and were analyzed using a mixed effects model with litter incorporated as a random effect, where post hoc tests were evaluated using a Bonferroni correction in R and data was plotted using GraphPad Prism 9. Statistical significance was defined as p<0.05.

#### Neuron morphology analysis quantification

Images of individual neurons were acquired as described above. Neuron morphology analysis was performed on single neurons. Within each litter, two embryos per sex/treatment were analyzed, with NSPCs from a single, independent embryo plated onto one coverslip (n=6 total embryos per sex/treatment from N=3 independent litters). From each embryo, five representative neurons were selected and analyzed from the same coverslip. For dendritic analyses, the number of primary, secondary, and tertiary dendrites was quantified for each neuron and averaged across the five neurons to obtain a representative value per slip/embryo. Dendritic length was calculated by first averaging the lengths of all dendrites for a given order (e.g., primary, secondary or tertiary) within each neuron (i.e., if multiple primary dendrites were present, these were averaged within the neuron) and then these values were averaged across the five neurons to obtain a representative value per slip/embryo. Total number of dendrites and cumulative dendritic length were calculated by summing all primary, secondary, and tertiary dendrites for all five neurons and averaged across the five neurons to obtain a representative value per slip/embryo. All measurements are reported as means ± SEM. Statistical significance was assessed using a mixed effects model with litter incorporated as a random effect, where post hoc tests were evaluated using a Bonferroni correction in R and data was plotted using GraphPad Prism 9. Statistical significance was defined as p<0.05.

#### QuPath quantification

Images of coronal brain sections were acquired as described above and analyzed using QuPath software. For all tissue-based counts and measurements, data are presented as means ± SEM. Each data point represents measurements obtained from a defined region of interest from a single embryo. Quantitative analyses were performed on tissue from n=3-4 embryos per sex/treatment derived from N=2-3 independent litters. Statistical analyses were performed using a mixed effects model with litter incorporated as a random effect, where post hoc tests were evaluated using a Bonferroni correction in R and data was plotted using GraphPad Prism 9. Statistical significance was defined as p<0.05.

## RESOURCE AVAILABILITY

### Lead contact

Requests for further information and resources and reagents should be directed to and will be fulfilled by the lead contact, Dr. Jessica M. Rosin.

### Materials availability

This study did not generate new unique reagents.

### Software availability

The Seurat R package (v5.2.1) was used for scRNAseq analysis, which is open-source and publicly available via CRAN and GitHub (Hao et al., 2024).

The QuPath software (v0.6.0) was used for FISH analysis, which is open-source and publicly available via GitHub (Bankhead et al., 2017).

### Data and code availability

All scRNAseq data from this study have been deposited in NCBI’s GEO (Edgar et al., 2002) and are accessible through GEO Series accession number GSE328919 (https://www.ncbi.nlm.nih.gov/geo/query/acc.cgi?acc=GSE328919).

All code used in this study is publicly available at https://github.com/RosinLabUBC/scRNAseq-code-KB.

All data reported in this paper will be shared by the lead contact upon request.

Any additional information required to reanalyze the data reported in this paper is available from the lead contact upon request.

## Notes

### Competing Interest Statement

The authors have declared no competing interest.

### Summary of Updates

All aspects of the manuscript (text, figures, supplemental material) have been revised.

