## Supplementary Figures 1-5 for "Prenatal exposure to maternal stress drives neurodevelopmental disruptions in the fetal hypothalamus"

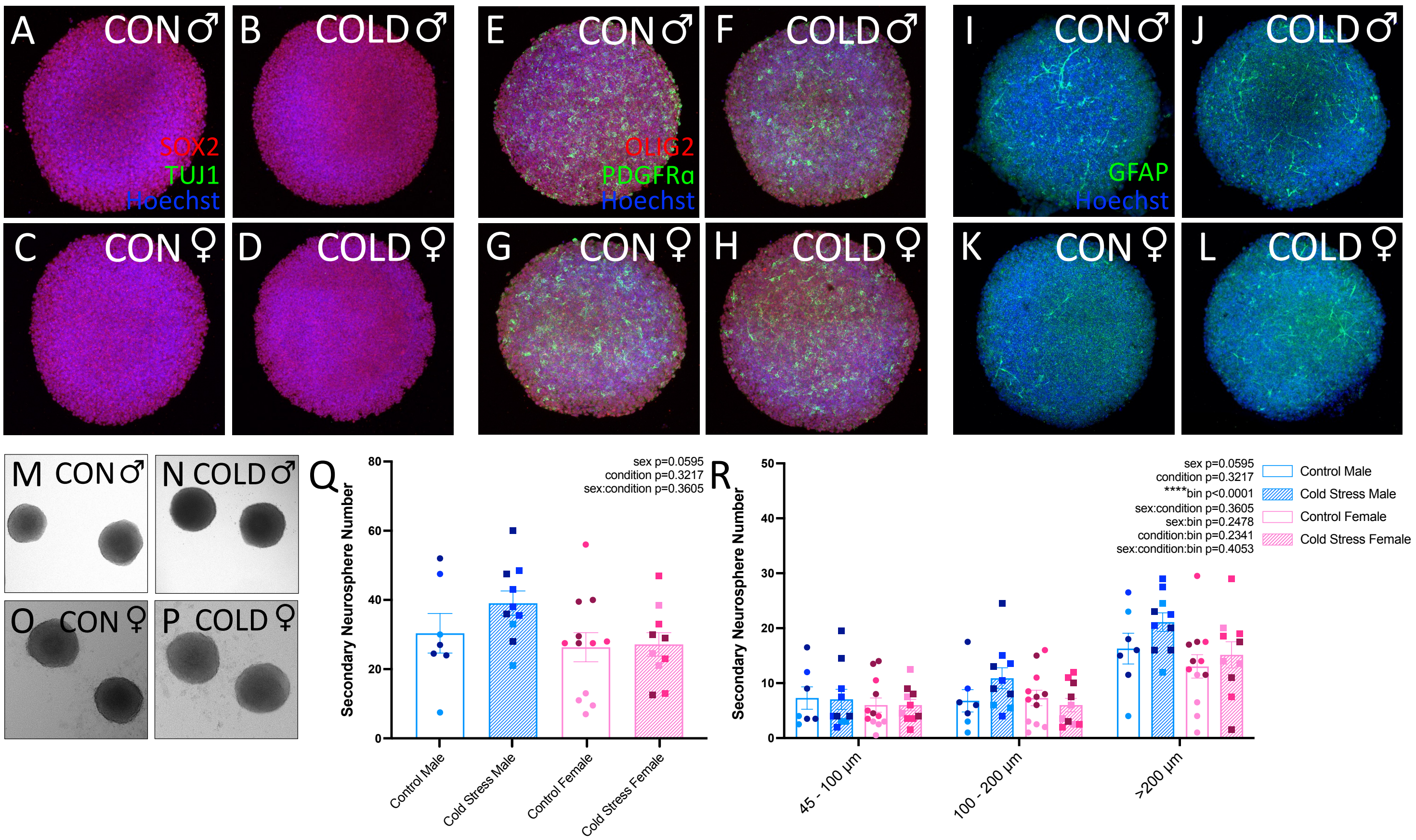

A CON♂
B COLD♂
SOX2
TUJ1
Hoechst
C CON♀
D COLD♀
E CON♂
F COLD♂
OLIG2
PDGFRα
Hoechst
G CON♀
H COLD♀
I CON♂
J COLD♂
GFAP
Hoechst
K CON♀
L COLD♀
M CON♂
N COLD♂
O CON♀
P COLD♀
Q
sex p=0.0595
condition p=0.3217
sex:condition p=0.3605
Secondary Neurosphere Number
Control Male
Cold Stress Male
Control Female
Cold Stress Female
R
sex p=0.0595
condition p=0.3217
****bin p<0.0001
sex:condition p=0.3605
sex:bin p=0.2478
condition:bin p=0.2341
sex:condition:bin p=0.4053
Control Male
Cold Stress Male
Control Female
Cold Stress Female
Secondary Neurosphere Number
45 - 100 µm
100 - 200 µm
>200 µm

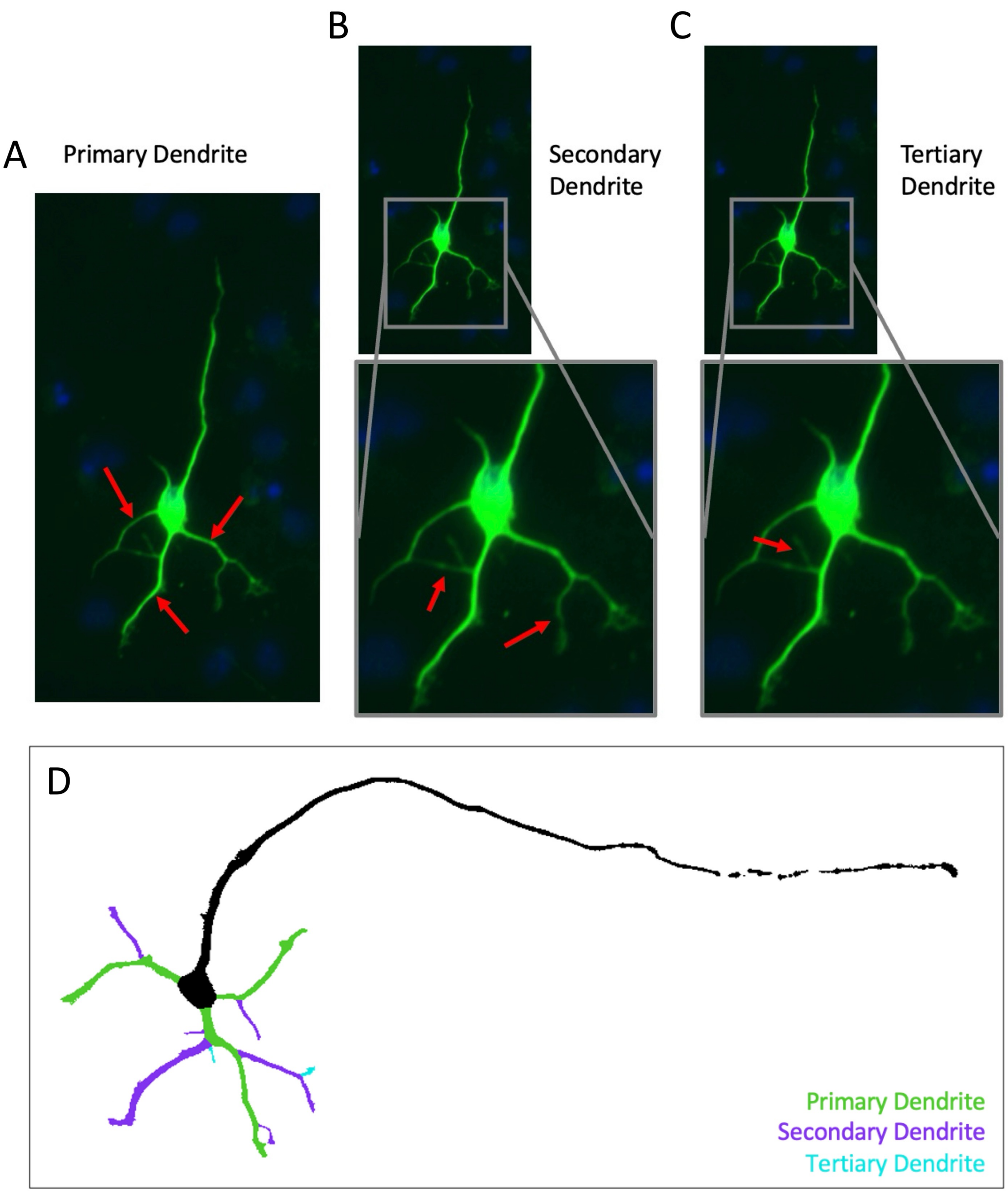
A
Primary Dendrite
B
Secondary
Dendrite
C
Tertiary
Dendrite
D
Primary Dendrite
Secondary Dendrite
Tertiary Dendrite

A

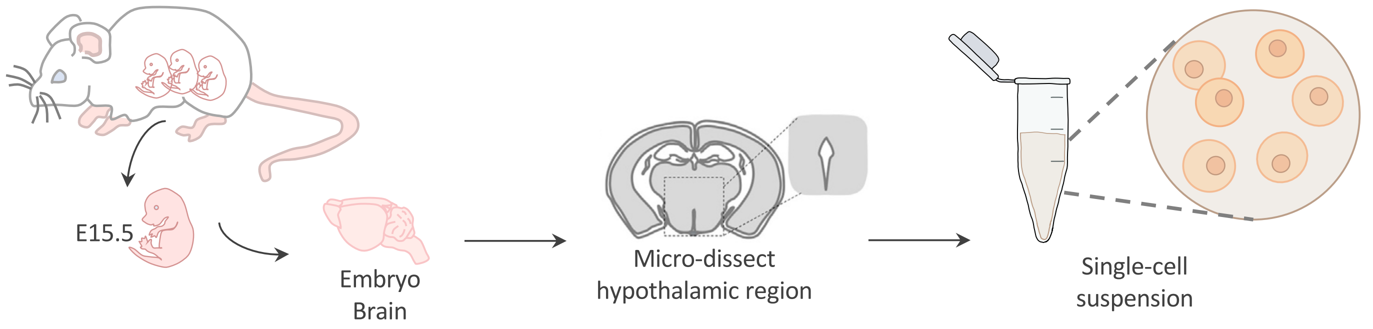


B

| Treatment Group | Control | Stress |
|---|---|---|
| Estimated Number of Cells | 15,556 | 17,961 |
| Mean Reads per Cell | 24,512 | 19,632 |
| Median UMI Counts per Cell | 7,406 | 7,240 |
| Median Genes per Cell | 3,097 | 3,101 |
| Total Number of Reads | 381,313,906 | 352,603,878 |
| Reads Mapped Confidently to Genome | 91.8% | 92.4% |
| Q30 bases in RNA Read | 95.5% | 92.1% |

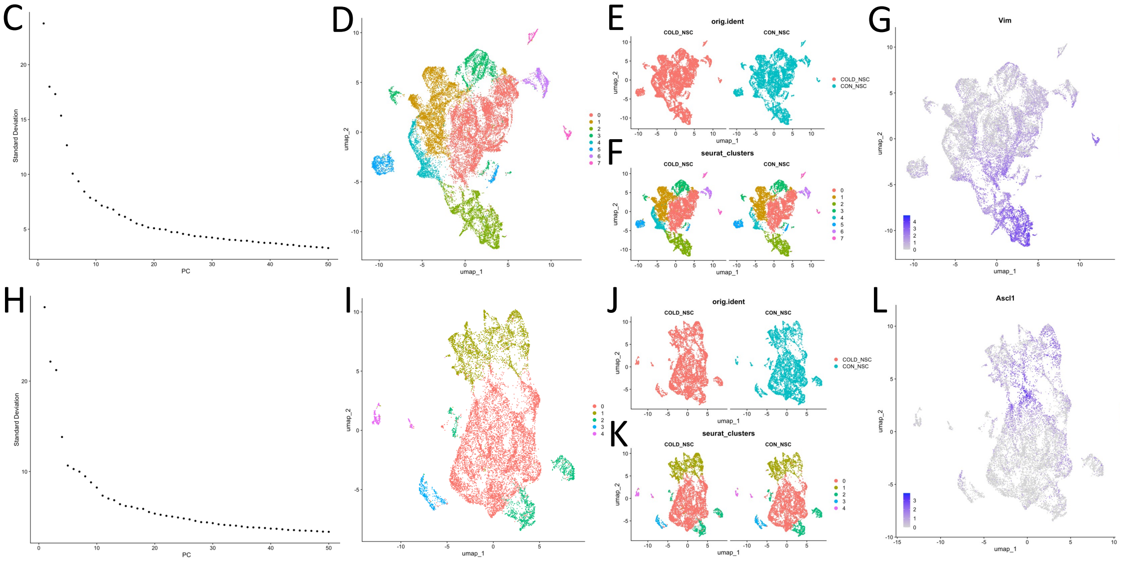


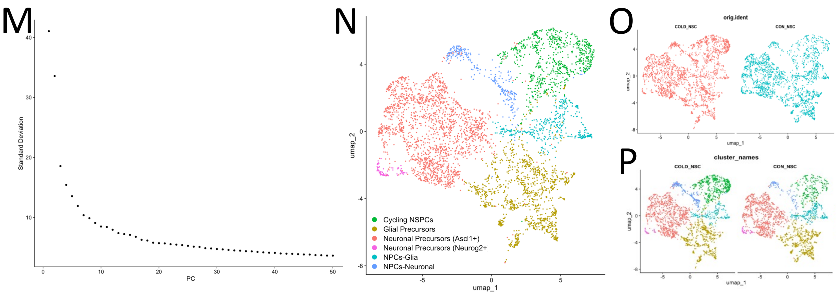


Q

| Treatment Group | Control | Stress |
|---|---|---|
| Number of Cells | 2,093 | 2,392 |
| Median UMI per Cell | 9,623 | 7,310 |
| Median Genes per Cell | 3,683 | 3,079 |

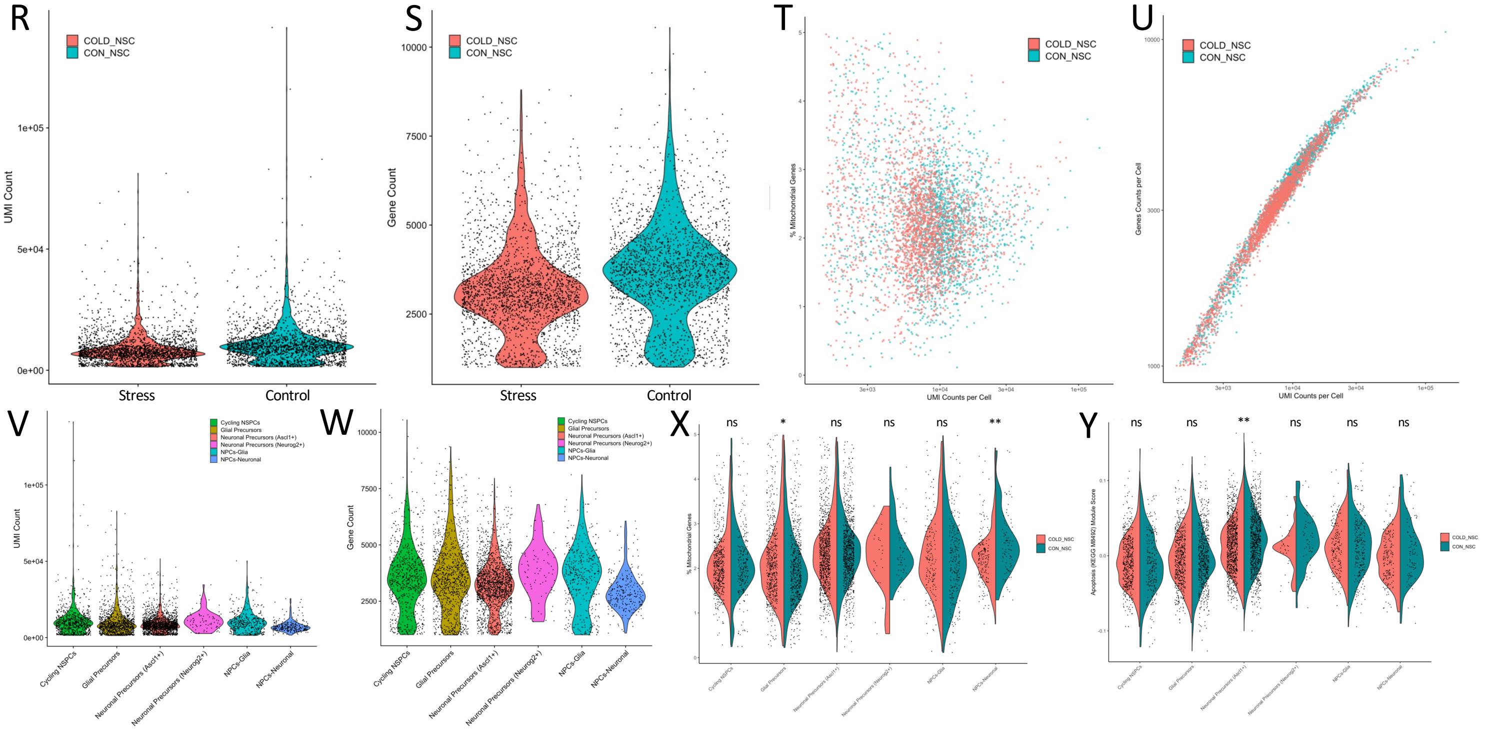

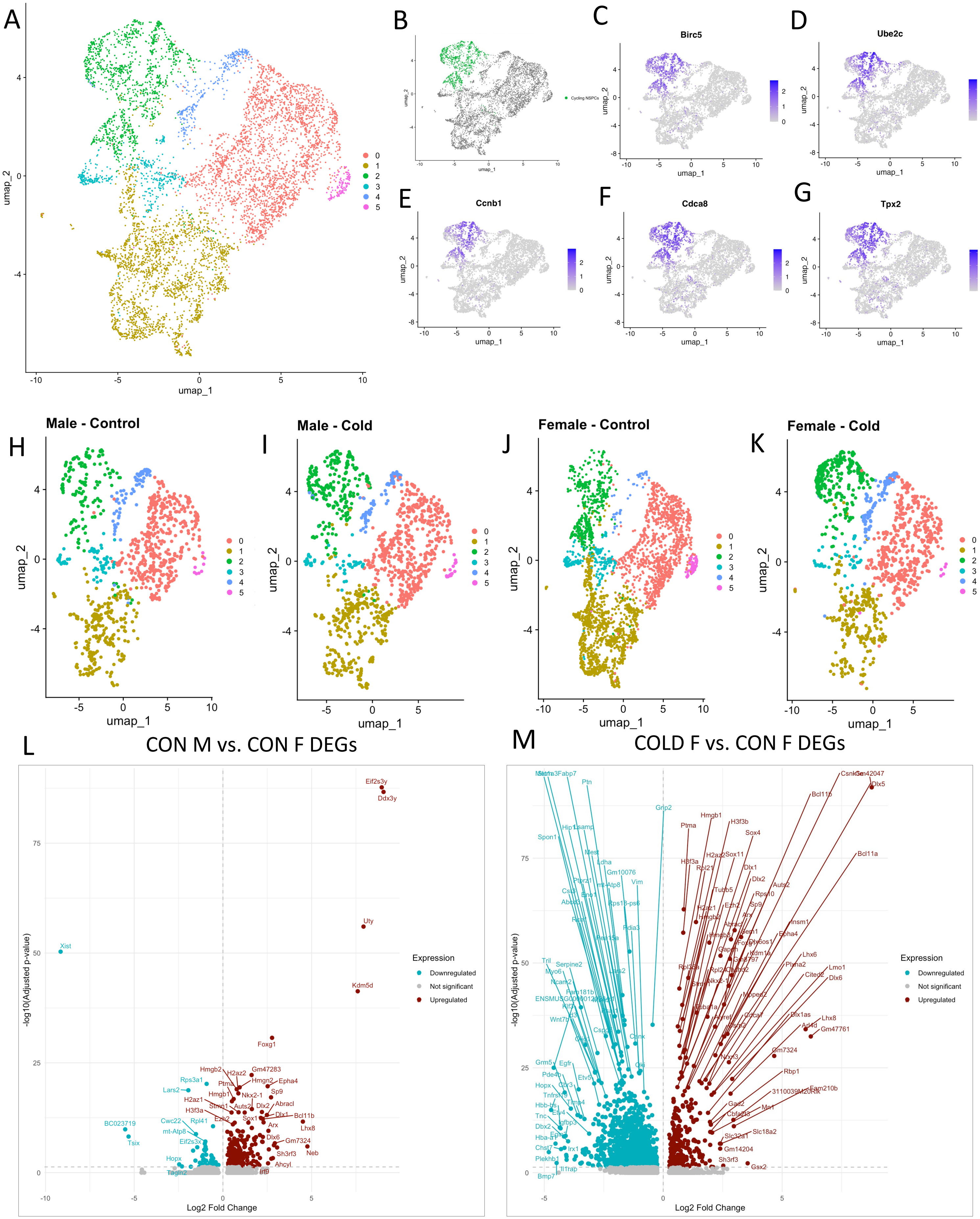

A
umap_1
umap_2
B
Cycling NSPCs
C
Birc5
D
Ube2c
E
Ccnb1
F
Cdca8
G
Tpx2
H
Male - Control
I
Male - Cold
J
Female - Control
K
Female - Cold
L
CON M vs. CON F DEGs
M
COLD F vs. CON F DEGs
Log2 Fold Change
-log10(Adjusted p-value)
Expression
Downregulated
Not significant
Upregulated
Eif2s3y
Ddx3y
Uty
Kdm5d
Xist
Foxg1
BC023719
Tsix
Csnk
Dlx5
Bcl11b
Bcl11a
Ptn
Grip2

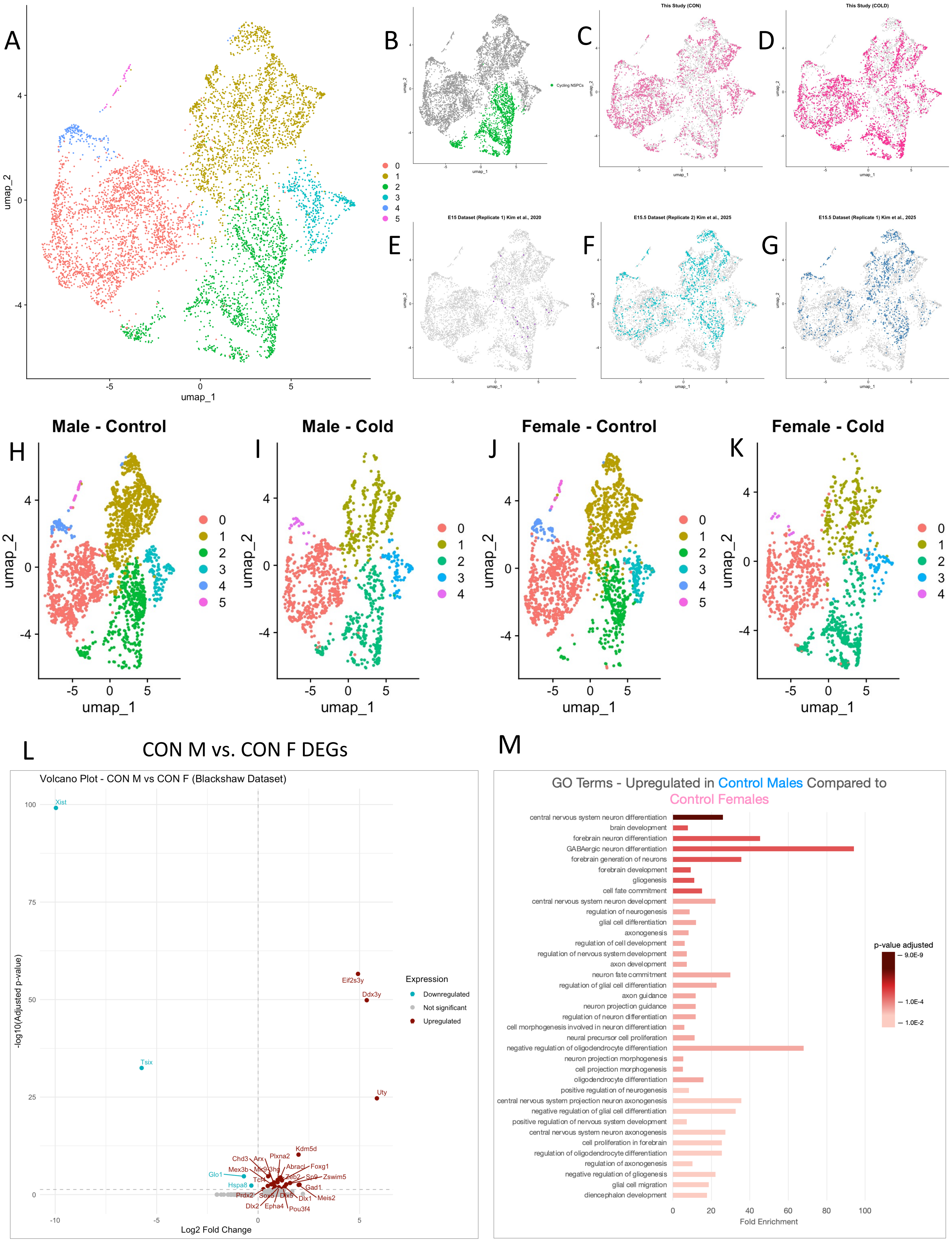

A
B
C
D
E
F
G
umap_1
umap_2
Cycling NSPCs
This Study (CON)
This Study (COLD)
E15 Dataset (Replicate 1) Kim et al., 2020
E15.5 Dataset (Replicate 2) Kim et al., 2025
E15.5 Dataset (Replicate 1) Kim et al., 2025
H
I
J
K
Male - Control
Male - Cold
Female - Control
Female - Cold
L
CON M vs. CON F DEGs
Volcano Plot - CON M vs CON F (Blackshaw Dataset)
-log10(Adjusted p-value)
Log2 Fold Change
Expression
Downregulated
Not significant
Upregulated
Xist
Tsix
Glo1
Hspa8
Eif2s3y
Ddx3y
Uty
Kdm5d
Chd3
Arx
Plxna2
Mex3b
Mir9-3hg
Abracl
Foxg1
Tcf4
Zeb2
Sp9
Zswim5
Gad1
Prdx2
Sox5
Dlx5
Dlx1
Meis2
Dlx2
Epha4
Pou3f4
M
GO Terms - Upregulated in Control Males Compared to Control Females
central nervous system neuron differentiation
brain development
forebrain neuron differentiation
GABAergic neuron differentiation
forebrain generation of neurons
forebrain development
gliogenesis
cell fate commitment
central nervous system neuron development
regulation of neurogenesis
glial cell differentiation
axonogenesis
regulation of cell development
regulation of nervous system development
axon development
neuron fate commitment
regulation of glial cell differentiation
axon guidance
neuron projection guidance
regulation of neuron differentiation
cell morphogenesis involved in neuron differentiation
neural precursor cell proliferation
negative regulation of oligodendrocyte differentiation
neuron projection morphogenesis
cell projection morphogenesis
oligodendrocyte differentiation
positive regulation of neurogenesis
central nervous system projection neuron axonogenesis
negative regulation of glial cell differentiation
positive regulation of nervous system development
central nervous system neuron axonogenesis
cell proliferation in forebrain
regulation of oligodendrocyte differentiation
regulation of axonogenesis
negative regulation of gliogenesis
glial cell migration
diencephalon development
Fold Enrichment
p-value adjusted
9.0E-9
1.0E-4
1.0E-2
