## Supplementary Figure & Table Legends for "Prenatal exposure to maternal stress drives neurodevelopmental disruptions in the fetal hypothalamus"

**Supplementary Figure 1. Maternal cold stress exposure does not impact the stem-like potential of E15.5 NSPCs.** (A-D) Secondary neurospheres stained with SOX2, TUJ1, and Hoechst. (E-H) Secondary neurospheres stained with OLIG2, PDGFRα, and Hoechst. (I-L) Secondary neurospheres stained with GFAP and Hoechst. (M-P) Representative images of secondary neurospheres generated from male and female NSPCs derived from primary neurospheres. (Q) Quantification of the number of secondary neurospheres generated from NSPCs derived from control male (n=7 embryos; N=3 litters), control female (n=12 embryos; N=3 litters), cold stress male (n=10 embryos; N=3 litters), and cold stress female (n=10 embryos; N=3 litters) primary neurospheres. (R) Quantification of the size distribution of secondary neurospheres generated from male and female NSPCs derived from control or cold stress primary neurospheres, which were binned by spheres sized between 45-100µm, 100-200µm or >200µm. Counts represent means ± SEM and were analyzed using a mixed effects model with litter incorporated as a random effect, where post hoc tests were evaluated using a Bonferroni correction. (Q, R) Each dot represents secondary neurospheres generated from primary neurospheres that were derived from a single, independent embryo. Dots of same color represent embryos from the same litter, while dots of different color represent embryos from different litters (N=3 independent litters).

**Supplementary Figure 2. Morphological analysis of neurons.** (A) Representative image of a neuron, where arrows point to the primary dendrites. (B) Representative image of a neuron, where red arrows in the high-magnification inset point to secondary dendrites. (C) Representative image of a neuron, where a red arrow in the high-magnification inset points to a tertiary dendrite. (D) Representative binary image of a neuron with annotated dendritic morphology.

**Supplementary Figure 3. Single-cell RNA sequencing metrics and quality controls.** (A) Schematic diagram illustrating the generation single-cell suspensions used for single-cell RNA sequencing experiments. (B) Raw sequencing metrics for each individual library (control vs cold stress) before subsetting. (C) Standard deviation for each of the first 50 principal components of cells collected from E15.5 control and cold stress male and female embryos pooled together (n=6; control: n=3 males, n=3 females; cold stress: n=4 males, n=2 females). (D) UMAP of all 22,643 cells collected from E15.5 control and cold stress male and female embryos. (E-F) UMAPs showing the distribution of cell clusters across control and cold stress treatment groups. (G) UMAP highlighting expression of *Vim*. (H) Standard deviation for each of the first 50 principal components of *Vim*+ cells collected from E15.5 control and cold stress male and female embryos. (I) UMAP of 13,624 *Vim*+ cells collected from E15.5 control and cold stress male and female embryos. (J-K) UMAPs showing the distribution of cell clusters across control and cold stress treatment groups. (L) UMAP highlighting expression of *Ascl1* in *Vim*+ cells. (M) Standard deviation for each of the first 50 principal components for cluster 1 and *Ascl1*+ cells collected from E15.5 control and cold stress male and female *Vim*+ cells. (N) UMAP for 4,485 cluster 1 and *Ascl1*+ cells collected from E15.5 control and cold stress male and female *Vim*+ cells, removing double positive and non-expressing cells that were assessed for male- (*Ddx3y, Uty, Eif2s3y*) and female-specific (*Xist*) transcripts. (O-P) UMAPs showing the distribution of cell clusters across control and cold stress treatment groups. (Q) Sequencing metrics for each individual library (control vs cold stress) after subsetting. (R) UMI counts. (S) Gene counts. (T)

Correlation of UMI counts with percent mitochondrial genes. (U) Correlation of UMI counts with gene counts per cell. (V) UMI counts grouped by cell cluster. (W) Gene counts grouped by cell cluster. (X) Violin plot depicting percent mitochondrial genes grouped by cell cluster (cycling NSPCs, p=0.50; glial precursors, p=0.042; neuronal precursors (*Ascl1+*), p=0.38; neuronal precursors (*Neurog2+*), p=0.38; NPCs-glia, p=0.38; NPCs-neuronal, p=0.0067). (Y) Violin plot depicting gene set enrichment for apoptosis grouped by cell cluster (cycling NPCs, p=0.36; glial precursors, p=0.77; neuronal precursors (*Ascl1+*), p=0.0026; neuronal precursors (*Neurog2+*), p=0.24; NPCs-glia, p=0.77; NPCs-neuronal, p=0.17). (X, Y) Pairwise t-tests were used to compare control and cold stress conditions within each cluster, with p-values adjusted across clusters using the Benjamini-Hochberg FDR method. Statistical significance was defined as $p<0.05$.

**Supplementary Figure 4. Validating E15.5 hypothalamic single-cell RNA sequencing comparisons using an independent litter of control female embryos.** (A) UMAP plot of 6,615 cells collected from E15.5 control and cold stress male and female embryos pooled together (replicate 1 control: n=3 males; replicate 2 control: n=5 females; replicate 1 cold stress: n=4 males, n=2 females). (B) UMAP highlighting cycling NSPCs cluster. (C-G) Unique gene expression signatures for the cycling NSPC cluster. (H-K) Individual UMAPs showing the distribution of the cell clusters across each sex and treatment group. (L) Volcano plot depicting DEGs for E15.5 control male cycling NSPCs (run 1) compared to control female cycling NSPCs (run 2). (M) Volcano plot depicting DEGs for E15.5 cold stress female cycling NSPCs (replicate 1) compared to control female cycling NSPCs (replicate 2).

**Supplementary Figure 5. Validating hypothalamic single-cell RNA sequencing comparison through incorporation of publicly available E15 and E15.5 hypothalamic datasets.** (A) UMAP plot combining E15.5 control and cold stress hypothalamic cells generated in this study with E15 wild-type cells from Kim et al., 2020 (GEO, GSE132355: GSM3860738 replicate 1) and E15.5 wild-type cells from Kim et al., 2025 (Kim et al., 2025 GEO GSE284492: GSM8685159 replicate 1 and GSM8685160 replicate 2). (B) UMAP plot highlighting the cycling NSPC cluster. (C) UMAP plot highlighting E15.5 control cells generated in this study. (D) UMAP plot highlighting E15.5 cold stress cells generated in this study. (E) UMAP plot highlighting E15 wild-type cells from Kim et al., 2020 (replicate 1). (F) UMAP plot highlighting E15.5 wild-type cells from Kim et al., 2025 (replicate 2). (G) UMAP plot highlighting E15.5 wild-type cells from Kim et al., 2025 (replicate 1). (H-K) Individual UMAPs showing the distribution of the cell clusters across each sex and treatment group. (L) Volcano plot depicting DEGs for E15/E15.5 male cycling NSPCs compared to female cycling NSPCs. (M) GO ORA terms identified for E15/E15.5 male cycling NSPCs compared to female cycling NSPCs

**Supplementary Table 1. Mixed effects model using a Bonferroni correction for number and size distribution of primary and secondary neurospheres derived from E15.5 male and female NSPCs.**

**Supplementary Table 2. Mixed effects model using Bonferroni correction for neuron, OPC, oligodendrocyte, and astrocyte proportions on culture slips.**

**Supplementary Table 3. Mixed effects model using Bonferroni correction for primary, secondary, tertiary, and cumulative dendrite number and length for cultured neurons.**

**Supplementary Table 4. Significantly upregulated and downregulated DEGs in cycling NSPCs across sex (control male versus control female) and prenatal maternal cold stress conditions (maternal cold stress male versus control male; maternal cold stress female versus control female), including independent validation and GO ORA terms enriched.**

**Supplementary Table 5. Significantly upregulated and downregulated DEGs in cycling NSPCs across sex (male versus female) when analyzed using data from E15.5 hypothalamic cells generated in this study combined with E15 hypothalamic cells from Kim et al., 2020 and E15.5 hypothalamic cells from Kim et al., 2025, and GO ORA terms enriched.**

**Supplementary Table 6. Mixed effects model using Bonferroni correction for frequency distribution of *Dlx2*+ and *Dlx5*+ cells in male and female hypothalamic sections.**

**Supplementary Table 7. Significantly upregulated and downregulated DEGs in GABAergic cells across sex (control male versus control female) and prenatal maternal cold stress conditions (maternal cold stress male versus control male; maternal cold stress female versus control female), including GO ORA terms enriched.**
